# Omnitemporal choreographies of IP_3_R and all five STIM/Orai underlie the complexity of mammalian Ca^2+^ signaling

**DOI:** 10.1101/2020.10.04.325480

**Authors:** Scott M. Emrich, Ryan E. Yoast, Ping Xin, Vikas Arige, Larry E. Wagner, Nadine Hempel, Donald L. Gill, James Sneyd, David I. Yule, Mohamed Trebak

**Affiliations:** Department of Cellular and Molecular Physiology, the Pennsylvania State University College of Medicine; 500 University Dr. Hershey, PA, 17033 USA; Department of Pharmacology, the Pennsylvania State University College of Medicine; 500 University Dr. Hershey, PA, 17033 USA; Penn State Cancer Institute, the Pennsylvania State University College of Medicine; 500 University Dr. Hershey, PA, 17033 USA; Department of Pharmacology and Physiology, University of Rochester, Rochester, NY 14642 USA; Department of Mathematics, The University of Auckland, 38 Princes Street, Auckland, 1010, New Zealand

## Abstract

Invertebrates express one endoplasmic reticulum (ER)-resident Ca^2+^-sensing stromal-interaction molecule (Stim) and one Orai plasma membrane channel protein. Stim conveys store depletion to Orai, mediating the evolutionarily conserved Ca^2+^ release-activated Ca^2+^ (CRAC) current. The crucial role of their vertebrate homologues, STIM1 and Orai1 in mediating CRAC activity in mammals is well-established. However, mammals possess two STIM and three Orai isoforms and the choreography of their interactions under physiological receptor activation is unknown. We show that the five mammalian STIM1/2 and Orai1/2/3 isoforms have non-redundant functions. Yet, all five isoforms are always required together to ensure the graded diversity of mammalian Ca^2+^ signaling events in response to the full spectrum of agonist strengths. Receptor-activated Ca^2+^ signaling across the range of stimulus intensities requires functional interactions between not only STIM1/2 and Orai1/2/3, but also IP_3_R, ensuring that receptor-mediated Ca^2+^ release is precisely tailored to Ca^2+^ entry and activation of nuclear factor of activated T-cells (NFAT). This is orchestrated by two interdependent and counterbalancing paradigms: the N-termini Ca^2+^-binding ER-luminal domains of *unactivated* STIM1/2 inhibit IP_3_R-evoked Ca^2+^ release. Gradual increase in agonist intensity leads to gradual STIM1/2 activation and relief of IP_3_R inhibition. Concomitantly, the cytosolic C-termini of *activated* STIM1/2 differentially interact with Orai1/2/3 proteins as agonist intensity increases. Thus, coordinated and omnitemporal functions of all five STIM/Orai proteins and IP_3_Rs at the ER-lumen and cytosol translate the strength of agonist stimulation to precise levels of Ca^2+^ release, Ca^2+^ entry and NFAT induction, ensuring the diversity and fidelity of complex mammalian Ca^2+^ signaling.

**Highlights:** - All five STIM/Orai and IP_3_R are always required together in mammalian Ca^2+^ signalling
- Unactivated STIM1/2 inhibit IP_3_R and activated STIM1/2 cooperatively activate Orai1/2/3
- STIM1 contribution increases and that of STIM2 decreases as agonist intensifies
- Graded IP_3_R disinhibition and Orai activation tailor receptor activity to NFAT induction

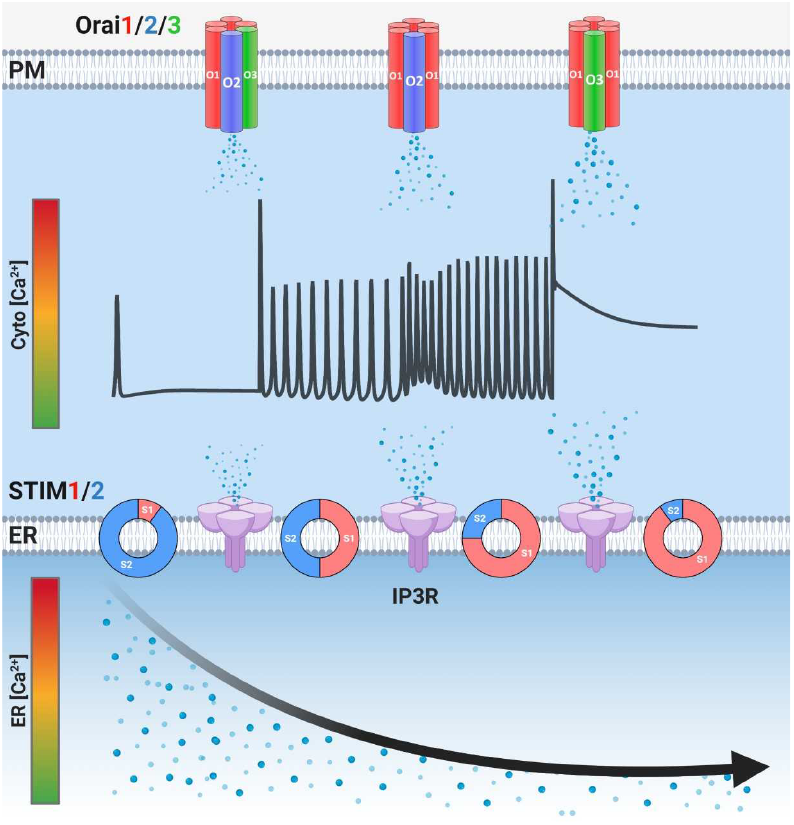

## Introduction

Ca^2+^ is a universal second messenger which regulates a myriad of physiological responses, including immune cell function, exocrine secretion and muscle contraction^1,2^. The endoplasmic reticulum (ER) luminal free Ca^2+^ is approximately ten thousand-fold higher than the cytosol^3^. This massive ER free Ca^2+^ content is tightly controlled through the opposing functions of the inositol 1,4,5-trisphosphate receptor (IP_3_R), the channel which releases ER Ca^2+^ into the cytosol and the sarcoplasmic/ER Ca^2+-^ATPase (SERCA), which pumps Ca^2+^ from the cytosol into the ER lumen^2-4^. Agonist stimulation of plasma membrane (PM) receptors coupled to phospholipase C (PLC) produces diffusible IP_3_, leading to activation of IP_3_Rs and depletion of ER Ca^2+^ stores^5^. Store depletion triggers the activation of ER resident stromal-interaction molecule (STIM) proteins to couple with and gate PM Orai channels and drive Ca^2+^ entry through the ubiquitous store-operated Ca^2+^ entry (SOCE) pathway^6-16^, mediated by the Ca^2+^ release-activated Ca^2+^ (CRAC) current^17^. SOCE plays a critical dual role, first in the acute refilling of ER Ca^2+^ stores to support Ca^2+^ release signals, and second to generate local Ca^2+^ signals that activate Ca^2+^-dependent nuclear factor of activated T-cells (NFAT) transcription factors^8,18^. Dysregulation of SOCE contributes to the pathogenesis of multiple immune, muscle, exocrine and metabolic disorders^14,19-22^.

Most protocols used to study SOCE use either SERCA blockers (e.g. thapsigargin) or high agonist concentrations, both of which induce massive non-physiological depletion of ER Ca^2+^ and robust cytosolic Ca^2+^ plateaus. Under these conditions, STIM1 and Orai1 have emerged as the major participants in SOCE. However, maximal activation of SOCE represents only the extreme end of the diverse and graded array of SOCE activities in response to the full range of agonist concentrations. Stimulation of mammalian cells with relatively low, physiological concentrations of agonists results in the generation of repetitive cytosolic Ca^2+^ oscillations^23^. In lymphocytes, Ca^2+^ oscillations are driven by oscillating SOCE activity independently of IP_3_Rs^24,25^. In most cells however, Ca^2+^ oscillations are initiated by IP_3_R-mediated Ca^2+^ release and maintained by ER Ca^2+^ store refilling through SOCE^26,27^. We recently demonstrated that Orai1 is dispensable for sustaining regenerative Ca^2+^ oscillations in HEK293 cells^28^. Surprisingly, endogenous Orai2 and Orai3, which contribute almost undetectable fraction of SOCE were able to maintain oscillations, suggesting that miniscule levels of SOCE are enough to support Ca^2+^ oscillations over extended periods of time^28^.

Although structurally similar, mammalian STIM1 and STIM2 are activated by distinct levels of ER Ca^2+^ depletion^12,13,29,30^. By comparison to STIM1, STIM2 is a weak activator of Orai channels and is triggered by more modest levels of ER Ca^2+^ depletion^29,31,32^. The contributions and precise choreographies of native STIM1 and STIM2 translocation to ER-PM junctions and interactions with Orai1/2/3 over the range physiological agonist concentrations is unknown. A number of studies have used gene knockdown approaches to investigate each STIM homolog in the regulation of Ca^2+^ oscillations^33-35^, but with conflicting results. Earlier reports suggested that STIM1 is solely required for Ca^2+^ oscillations in response to low agonist concentrations^35,36^. Conversely, other studies suggested that STIM2 is the key regulator of Ca^2+^ oscillations^33,34^. Further complicating this picture, evidence has emerged that STIM2 actively recruits STIM1 to ER-PM junctions under conditions of modest ER Ca^2+^ depletion^33,37,38^, suggesting that STIM1/2 play a dynamic and cooperative role in Ca^2+^ signaling than previously appreciated.

Here, we generated single and double STIM1/STIM2 knockout HEK293 cells using CRISPR/Cas9 gene-editing and double Orai/single STIM knockout cells (i.e. cells expressing one native STIM and one native Orai) to elucidate how each native STIM communicates with each native Orai to shape cytosolic Ca^2+^ profiles. We show that both STIM isoforms are always required together to ensure the fidelity of Ca^2+^ signaling events in response to different agonist concentrations. The distinct C-termini of STIM1 and STIM2 coordinate the wide range of Ca^2+^ oscillation frequencies in response to low-and mid-range agonist concentrations, while the C-terminus of STIM1 exclusively drives sustained Ca^2+^ plateaus in response to high concentrations of agonist. Cooperation between STIM1 and STIM2 is required for driving optimal nuclear translocation of NFAT1 and NFAT4 under both low and high agonist concentrations. We demonstrate that the Ca^2+^ binding N-terminal EF-hand of unactivated (resting) STIMs are required for STIM1/2-mediated negative regulation of IP_3_Rs. Mathematical modeling revealed STIM1/STIM2 cooperation and regulation of IP_3_Rs by unactivated STIM proteins. The model predicted that under basal conditions, 93% of SOCE activity is mediated by STIM2 and the remainder is by STIM1. However, the contribution of STIM2 to SOCE decreases while that of STIM1 increases with increasing agonist concentrations and responses in the form of Ca^2+^ plateaus are exclusively mediated by STIM1.

Our data challenge the current oversimplified model of mammalian CRAC channel activation involving either STIM2/Orai1 or STIM1/Orai1 interactions under basal and agonist stimulation, respectively. Physiological CRAC channel activities are *Omni tempore* choreographies of all five STIM/Orai proteins and their functional interaction with IP_3_Rs. Unactivated STIMs inhibit IP_3_Rs, likely setting a threshold for IP_3_ activation of IP_3_Rs. As the ratio of activated/unactivated STIMs increases with increasing agonist concentrations, IP_3_R inhibition by STIMs is relieved concomitantly with increased CRAC activity, effectively matching Ca^2+^ release to Ca^2+^ entry. While STIM1 and Orai1 alone can clearly generate robust Ca^2+^ signals as is the case for invertebrate Stim and Orai, the concomitant involvement of all five mammalian STIM/Orai isoforms and their interaction with IP_3_Rs are required for fine-tuning the diverse and graded Ca^2+^ signals and NFAT activities in response to the diverse physiological agonist intensities and types encountered by complex mammalian cells.

## Results

### Both native STIM1 and STIM2 support Ca^2+^ oscillations

We generated several clones of HEK293 cells lacking STIM1 (STIM1-KO), STIM2 (STIM2-KO), or both (STIM1/2-KO) using CRISPR/Cas9 (**Fig. 1A**). Western blot analysis demonstrated no compensatory change in protein expression of either STIM protein (**Fig. 1A**) or the three isoforms of IP_3_Rs (**S1. D-G**) in STIM knockout cells. There was no difference in Ca^2+^ release between parental wild-type (WT)-HEK293 and STIM-KO cells in response to store depletion induced by 2µM thapsigargin (**Fig. 1B;** see also **Fig. 4F** for ER measurements using CEPIA1*er*). As expected however, SOCE was slightly reduced in STIM2-KO cells, mostly reduced in STIM1-KO cells and completely abrogated in STIM1/2-KO cells (**Fig. 1B,C**). Orai1/2/3 triple knockout cells (Orai-TKO), which show complete abrogation of SOCE^28^, were included as controls. In our previous studies, we determined that 10 μM of the muscarinic receptor agonist carbachol (CCh) induces a sustained Ca^2+^ oscillatory response^28^. For each condition, five representative traces from individual cells are shown (**Fig. 1D-H**) and data from several independent experiments performed on two independent CRISPR/Cas9 KO clones per condition are statistically analyzed and reported (**Fig. 1I-L**). These data report for each condition the frequency of Ca^2+^ oscillations/cell, the percentage of oscillating cells, the percentage of cells responding with sustained Ca^2+^ plateaus and the percentage of non-responding cells (cells with either 0 or 1 spike) as defined in **S1. A-C**.

**Fig. 1.**
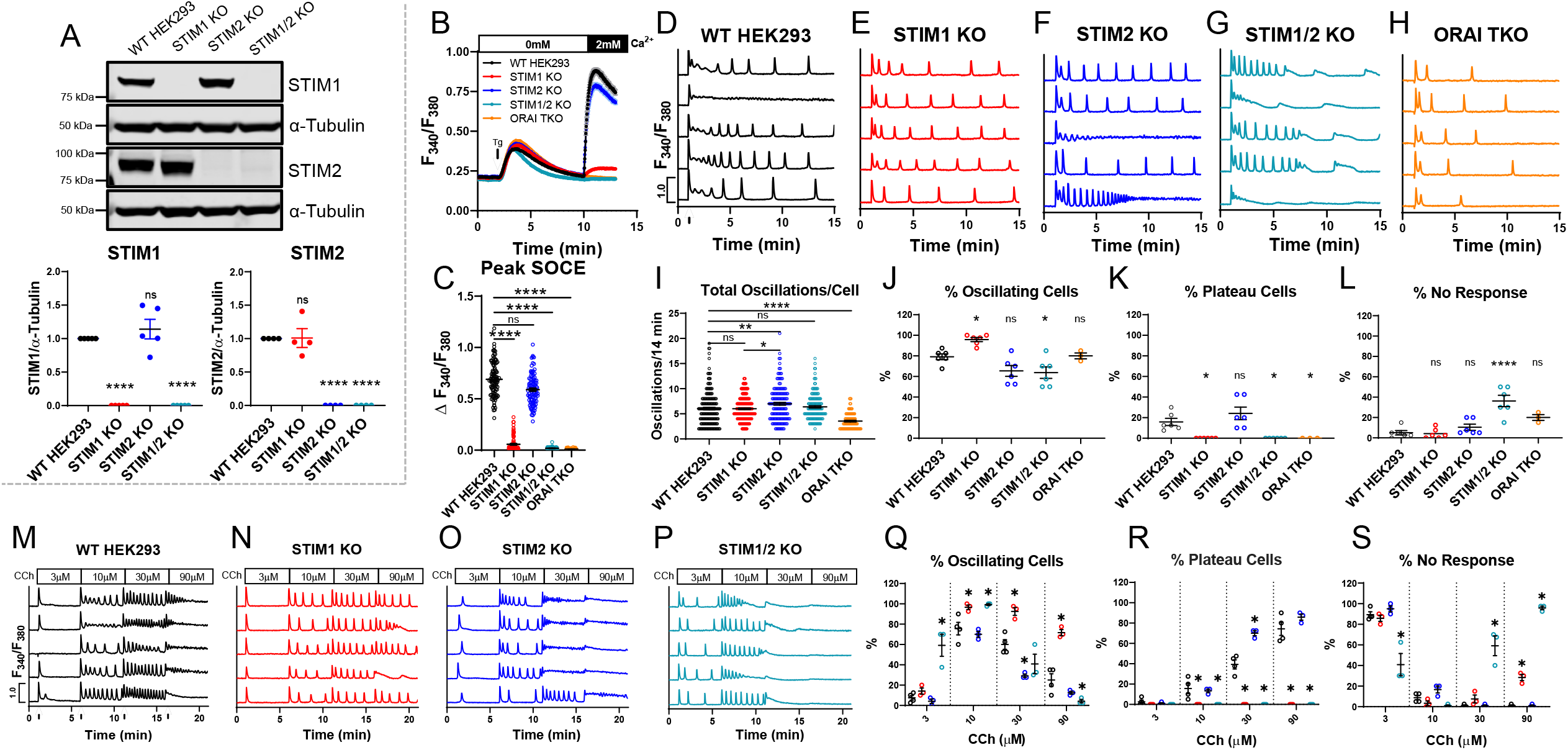
Native STIM1/STIM2 mediate unique Ca^2+^ oscillation profiles and agonist sensitivities. (A) *Top:* Western blot analysis of STIM1 and STIM2 proteins in WT-HEK293 and STIM knockout cells. *Bottom:* Quantification of STIM1 and STIM2 band intensity normalized to their respective α-tubulin loading control. Each point represents an individual biological replicate (*n* = 5 for STIM1, *n* = 4 for STIM2). (B) Ca^2+^ measurements with Fura2 upon store depletion with 2µM thapsigargin in 0mM Ca^2+^ followed by re-addition of 2mM Ca^2+^ to the external solution. (C) Quantification of maximal SOCE for all conditions in (B). From left to right *n* = 100, 101, 100, 99, and 120 individual cells. (D-H) Representative Ca^2+^ oscillation traces (Fura2) from 5 cells/condition upon stimulation with 10 µM carbachol (CCh) at 1 minute (indicated by arrow in D) in the presence of 2mM external Ca^2+^. (I) Quantification of total oscillations in 14 minutes. From left to right *n* = 189, 230, 156, 152, and 96 cells. (J) % oscillating cells, (K) % plateau cells, and (L) % non-responding cells for each condition in (D-H). For (J-L) from left to right *n* = 6, 6, 6, 6, and 3 independent experiments. (M-P) Representative Ca^2+^ traces in response to increasing CCh concentrations in the presence of 2mM Ca^2+^. Cells were sequentially stimulated with 3, 10, 30 and 90 µM CCh for 5 minutes each (indicated by arrows in M). (Q) Quantification of % oscillating cells, (R) % plateau cells, and (S) % non-responding cells for each condition in (M-P). For (Q-S) from left to right *n* = 4, 3, 3, and 3 independent experiments. All scatter plots of peak SOCE (C) and oscillation frequency (I) are presented as mean ± SEM and analyzed with the Kruskal-Wallis one-way ANOVA with multiple comparisons to WT-HEK293 cells where (*p<0.05; **p<0.01; ****p<0.0001; ns, not significant). Scatter plots in (A, J-L, Q-S) are presented as mean ± SEM and analyzed with one-way ANOVA with multiple comparisons to WT-HEK293 cells where (*p<0.05; ****p<0.0001).

**Fig. 4.**
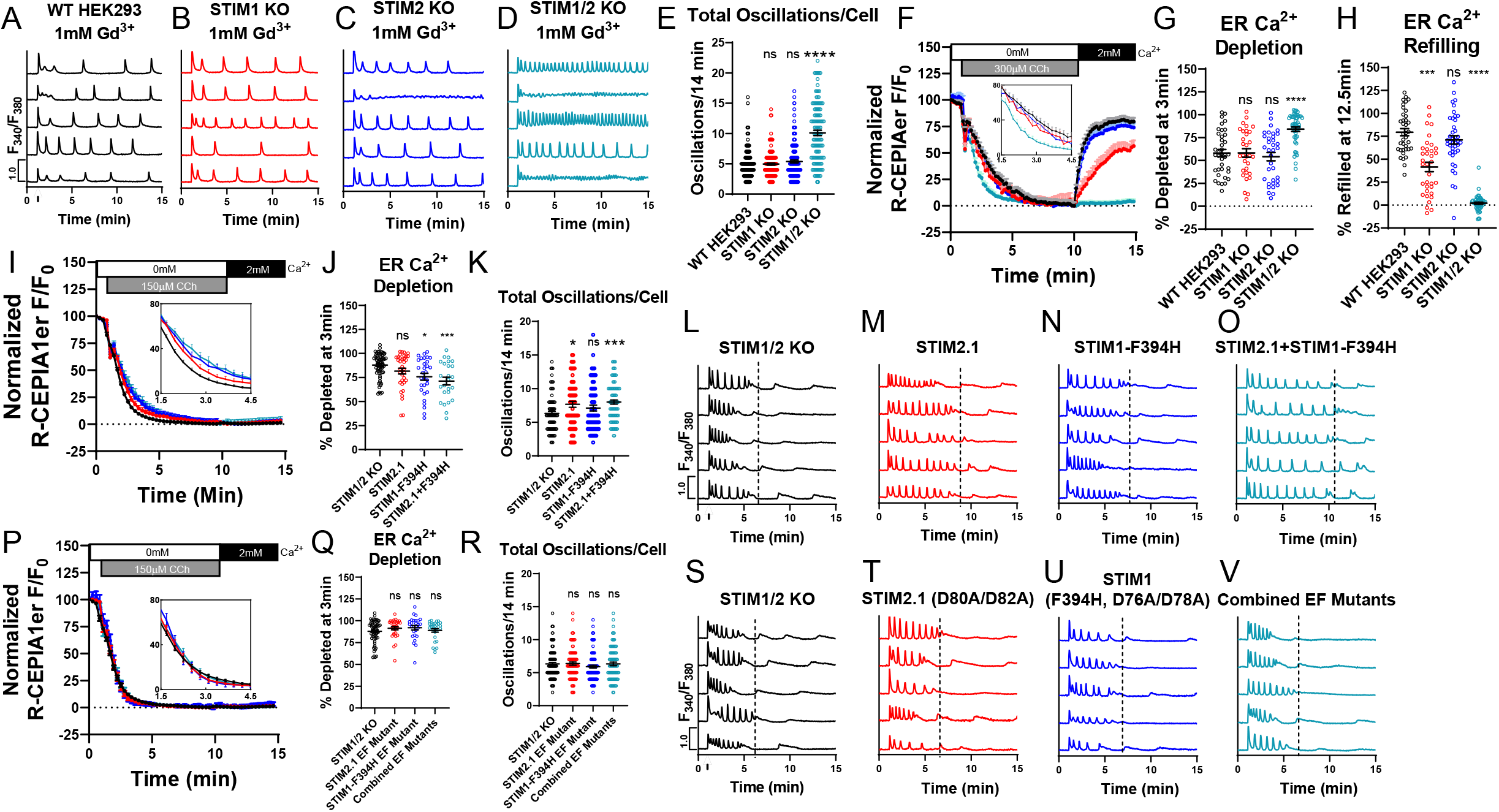
N-terminal EF-hands of STIM1/2 regulate receptor-mediated Ca^2+^ release. (A-D) Representative Ca^2+^ traces from 5 cells/condition with 10 µM CCh stimulation at 1 minute (arrow in A) with 1mM Gd^3+^ and 2mM Ca^2+^ in bath. (E) Total oscillations/14 minutes from (A-D). Left to right, *n*=173, 191, 166, and 123 cells. (F) ER Ca^2+^ measurements in cells using R-CEPIA1*er*. Cells were stimulated with 300µM CCh in 0mM Ca^2+^ followed by re-addition of 2mM Ca^2+^ at 10 minutes (without CCh). Inset: zoomed-in from 1.5-4.5 minutes. (G) ER Ca^2+^ depletion rate in (F) at 3 minutes after CCh stimulation in 0mM Ca^2+^. (H) ER Ca^2+^ refilling rate in (F) at 12.5 minutes in 2mM Ca^2+^. Left to right *n*=37, 35, 41, and 53 cells. (I) ER Ca^2+^ measurements in STIM1/2-KO or STIM1/2-KO cells rescued with YFP-STIM2.1, STIM1-F394H-CFP, or STIM2.1 + STIM1-F394H. Inset: zoomed-in from 1.5-4.5 minutes. (J) ER Ca^2+^ depletion rate in (I) at 3 minutes after CCh stimulation in 0mM Ca^2+^. Left to right *n*=67, 36, 30, and 25 cells. (K) Total oscillations/14 minutes from STIM1/2-KO or STIM1/2-KO cells individually rescued with YFP-STIM2.1, STIM1-F394H-CFP, or combined STIM2.1 + STIM1-F394H. Cells were stimulated with 10 µM CCh in 2 mM Ca^2+^. Left to right *n*=81, 64, 67, and 88 cells. (L-O) Representative Ca^2+^ traces from 5 cells/condition from (K). (P) ER Ca^2+^ measurements in STIM1/2-KO or STIM1/2-KO cells rescued with YFP-STIM2.1-(D80A/D82A), STIM1-(F394H/D76A/D78A)-CFP, or STIM2.1+STIM1-F394H EF-hand mutants. Inset: zoomed-in from 1.5-4.5 minutes. (Q) ER Ca^2+^ depletion rate in (P) at 3 minutes after CCh stimulation in 0mM Ca^2+^. Left to right *n*=67, 31, 26, and 29 cells. (R) Total oscillations/14 minutes from (Q) after stimulation with 10 µM CCh in 2mM Ca^2+^. Left to right *n*=108, 97, 101, and 93 cells. (S-V) Ca^2+^ traces from 5 cells/condition from (R). Scatter plots of oscillation frequency (E, K, R), ER Ca^2+^ depletion (G, J, Q), and ER Ca^2+^ refilling (H) as mean ± SEM analyzed with the Kruskal-Wallis one-way ANOVA with multiple comparisons to WT-HEK293 (E, G, H) or non-transfected STIM1/2-KO controls (J-K, Q-R) (*p<0.05; ***p<0.001; ****p<0.0001; ns, not significant).

Stimulation of WT-HEK293 cells with 10 μM CCh in the presence of 2mM extracellular Ca^2+^ produced an average of 6 oscillations/14 minutes (**Fig. 1I**). At this concentration, WT-HEK293 cells demonstrated an average of 79% oscillating cells, 16% plateau cells, and 5% non-responders (**Fig. 1J-L**). While STIM1-KO cells stimulated with 10 μM CCh demonstrated a similar oscillation frequency compared to WT-HEK293 (**Fig. 1I**), 95% of those cells responded with Ca^2+^ oscillations and 0% with sustained plateaus (**Fig. 1J-K**). STIM2-KO cells produced an increased average of ∼6.9 oscillations/14 minutes, as well as an increased percentage of plateau cells (24%) compared to WT-HEK293 (**Fig. J-L**). These results suggest that while both STIM1 and STIM2 can independently support Ca^2+^ oscillations, Ca^2+^ plateaus strictly require STIM1 and that STIM2 negatively regulates these plateau responses.

Unexpectedly, we observed rapid oscillations within the first 5 minutes of CCh stimulation in all oscillating STIM1/2-KO cells, followed by progressive decrease in peak amplitude and subsequent rundown (**Fig. 1G**). STIM1/2-KO cells showed a significantly increased percentage of non-responding cells (36%) and no cells eliciting a plateau response (**Fig. 1J-L**). The rapid oscillatory phenotype of STIM1/2-KO cells was unrelated to Ca^2+^ entry as inhibition of SOCE with 5µM Gd^3+^ reduced the frequency of Ca^2+^ oscillations in WT-HEK293 but not in STIM1/2-KO cells (**S2. A-E**). Furthermore, Orai-TKO cells have a significantly reduced frequency of oscillations with only ∼3 oscillations/14 min (**Fig. 1H-I** see also^28^). These findings suggest that STIM proteins provide an additional control mechanism over the rate of ER Ca^2+^ release through IP_3_Rs. This behavior of STIM1/2-KO cells will be addressed further below in the results section. Quantification of oscillation data from two independent CRISPR/Cas9 knockout clones per condition is presented in (**S3. A-H**).

Interestingly, we also observed a qualitative change in the behavior of Ca^2+^ oscillations between WT-HEK293 cells and STIM-KO cells as summarized in (**S1. H-K**). The oscillatory behavior of WT-HEK293 cells stimulated with 10µM CCh is typically rapid at first and gradually transitions into sustained oscillations where the descending phase of each oscillation fully returns to baseline before initiation of the next spike (**S1. H**). STIM2-KO cells show a reversal of this phenotype with oscillations that become rapid and tetanize into moderate plateaus (**S1. J**). STIM1-KO and STIM1/2-KO cells show consistent behavior throughout recordings (**S1. I, K**). Thus, while endogenous STIM1 and STIM2 can independently support Ca^2+^ oscillations, they likely function together under physiological conditions to fine-tune Ca^2+^ signaling. This concept is explored further below.

### STIM1/STIM2 cooperate over the range of agonist concentrations

Although STIM1 and STIM2 share a high degree of sequence homology, their ER-luminal N-terminal EF-hand domain of STIM2 displays a significantly lower affinity for calcium^32^. Several studies have proposed that basal SOCE activity (without agonist stimulation) is controlled by STIM2 while STIM1 proteins are increasingly recruited with increasing agonist concentrations^33,34^. We measured Ca^2+^ signals to increasing CCh concentrations ranging from 3-90 μM. At the lowest concentration of CCh (3 μM), WT-HEK293, STIM1-KO, and STIM2-KO cells were almost entirely refractory, while nearly 60% of STIM1/2-KO cells demonstrated Ca^2+^ oscillations (**Fig. 1M-S**). At 10 μM CCh, WT-HEK293 and STIM2-KO cells showed 75% and 70% of oscillating cells respectively, with 100% of STIM1-KO and STIM1/2-KO cells oscillating. Interestingly, STIM2-KO cells demonstrated an increased propensity to respond by exhibiting plateaus at 30 μM CCh (70%) compared to WT-HEK293 cells (40%), suggesting that STIM2 also negatively regulates STIM1 and SOCE at low CCh concentrations (3-10 µM). At the highest CCh concentration (90 µM), WT-HEK293 and STIM2-KO cells reached comparable percentages of cells responding with plateaus (∼80%). Importantly, STIM1-KO cells failed to support plateaus at all CCh concentrations (**Fig. 1Q-S**), further documenting that plateaus are exclusively mediated by STIM1. These data suggest that STIM1 and STIM2 work collaboratively across the range of stimulus intensities to tailor Ca^2+^ responses to their respective agonist concentrations.

### STIM C-termini determine the diversity of receptor-activated Ca^2+^ signaling events

We utilized STIM1/2-KO cells to determine whether STIM1 and STIM2 re-expression can rescue the phenotype of STIM1/2-KO cells. In order to rescue STIM proteins to near endogenous protein levels, we expressed YFP-tagged STIM constructs driven by the relatively weak thymidine kinase (tk) promoter^39^ (**S4. A** shows Western Blots documenting re-expression of STIM1/2 to near endogenous levels). Both tk-STIM1 and tk-STIM2 constructs rescued SOCE activated by store depletion with 2 µM thapsigargin. As expected, tk-STIM1 supported a significantly bigger peak SOCE compared to tk-STIM2 (**Fig. 2F, H**). Furthermore, 10 µM CCh induced sustained oscillations and plateaus in STIM1/2-KO cells expressing tk-STIM1 with similar percentages to those of STIM2-KO cells (**Fig. 2A-C; J-L**). Similarly, tk-STIM2 expression in STIM1/2-KO cells supported oscillations with a percentage similar to that of STIM1-KO cells, with 0% of cells displaying plateaus (**Fig. 2A-C, J-L**). Frequencies of oscillations were generally higher in these rescue experiments with tk-STIM1 and tk-STIM2 supporting ∼12.6 and 9 oscillations/14 min, respectively (**Fig. 2I**), suggesting that while average tk-STIM1 and tk-STIM2 protein expression within cell populations is similar to endogenous STIM levels (**S4. A**), tk-STIM1 and tk-STIM2 protein levels in individual cells are likely higher compared to native expression.

**Fig. 2.**
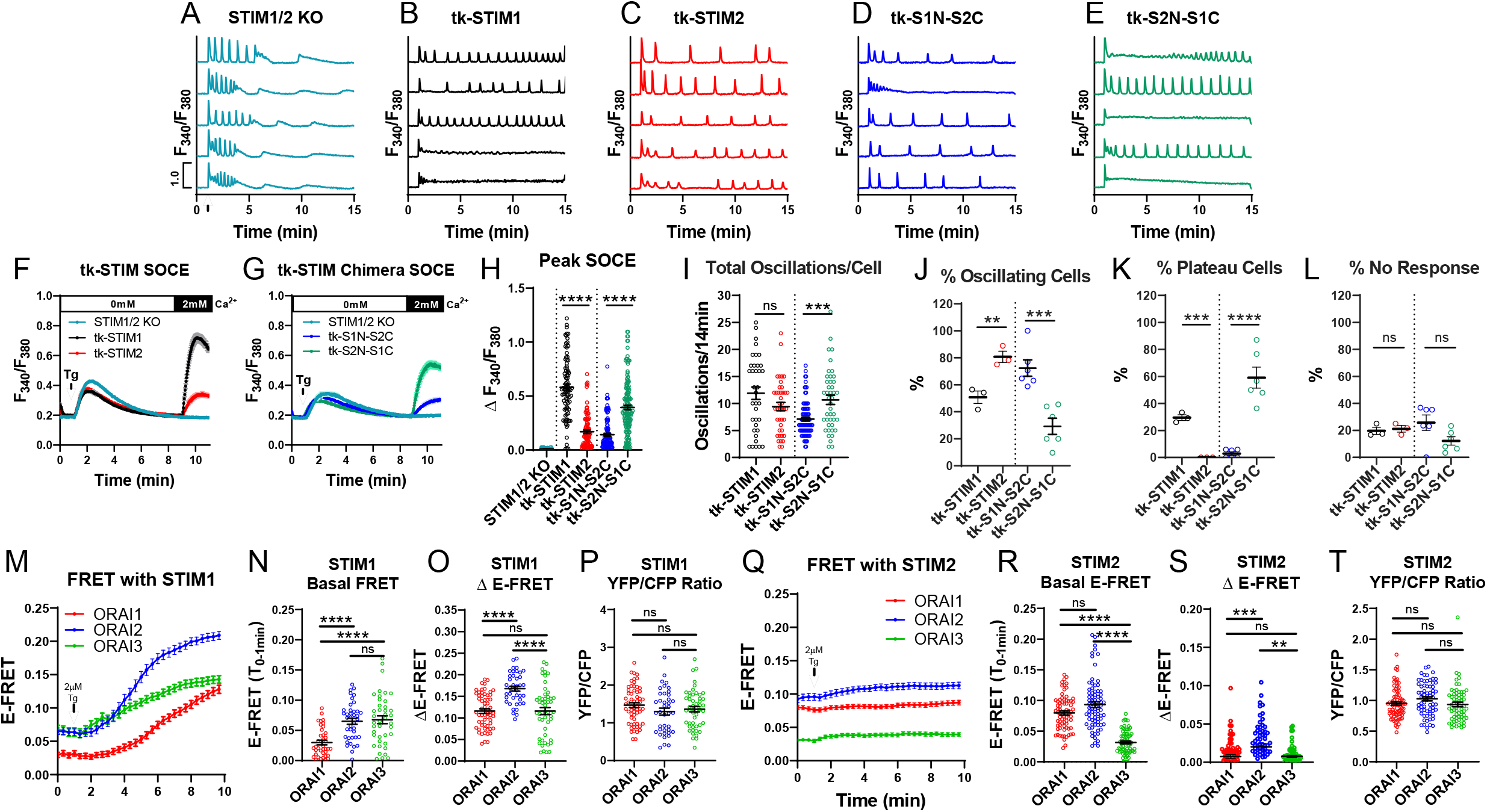
STIM1/STIM2 have distinct interactions with Orai1/2/3. (A-E) Ca^2+^ traces of (A) non-transfected STIM1/2-KO or STIM1/2-KO cells individually rescued with (B) tk-YFP-STIM1, (C) tk-YFP-STIM2, (D) tk-YFP-S1N-S2C, and (E) tk-YFP-S2N-S1C from 5 representative cells/condition upon stimulation with 10 µM CCh at 1 minute (arrow in A) in the presence of 2mM Ca^2+^. (F) SOCE in response to 2µM thapsigargin in Ca^2+^-free followed by re-addition of 2mM Ca^2+^ in STIM1/2-KO or STIM1/2-KO cells rescued with tk-YFP-STIM1 or tk-YFP-STIM2. (G) SOCE in STIM1/2-KO cells rescued with tk-YFP-S1N-S2C or tk-YFP-S2N-S1C. (H) Maximal SOCE for (F-G). From left to right *n* = 90, 88, 96, 116, and 154 cells. (I) Total oscillations in 14 minutes for (A-E). From left to right *n* = 36, 41, 87, and 43 cells. (J) % of oscillating, (K) plateau, and (L) non-responding cells for (B-E); from left to right *n* = 3, 3, 6, and 6 independent experiments. (M) E-FRET plotted as mean ± SEM between STIM1-YFP and each CFP-Orai expressed in Orai-TKO cells at baseline and after 2 µM thapsigargin in 2mM Ca^2+^. (N) Scatter blots quantifying basal E-FRET, (O) change of E-FRET after thapsigargin, and (P) YFP/CFP fluorescence ratios from data in (M). (N-P) from left to right n = 60 (Orai1), 40 (Orai2), and 51 (Orai3) cells. (Q) E-FRET presented as mean ± SEM between STIM2-YFP and each CFP-Orai expressed in Orai-TKO cells before and after thapsigargin. (R) Scatter blots quantifying basal E-FRET, (S) change of E-FRET after thapsigargin, and (T) YFP/CFP fluorescence ratios from data in (Q). (R-T) from left to right *n* = 71 (Orai1), 67 (Orai2), and 64 (Orai3) cells. Scatter plots of peak SOCE (H) and oscillation frequency (I) are presented as mean ± SEM and analyzed with the Mann-Whitney U test (***p<0.001; ****p<0.0001; ns, not significant). Scatter plots in (J-L) are presented as mean ± SEM and analyzed with an unpaired Student’s t-test (**p<0.01; ***p<0.001; ****p<0.0001; ns, not significant). Scatter plots of E-FRET quantifications (N-P, R-T) are plotted as mean ± SEM and analyzed with the Kruskal-Wallis one-way ANOVA with multiple comparisons (*p<0.05; **p<0.01; ****p<0.0001; ns, not significant).

Next, we determined the functional significance of the N-terminal versus the C-terminal domains of each STIM isoform in regulating Ca^2+^ oscillations by utilizing two chimeric constructs that we recently developed^40^. The first chimera consists of the N-terminal and transmembrane domains of STIM1 combined with the C-terminus of STIM2 (tk-S1N-S2C) while the second consists of the N-terminal and transmembrane domains of STIM2 combined with the C-terminus of STIM1 (tk-S2N-S1C). Both STIM chimeras rescued SOCE triggered by thapsigargin when expressed in STIM1/2-KO cells with tk-S2N-S1C demonstrating significantly higher Ca^2+^ entry compared to tk-S1N-S2C (**Fig. 2G-H**). Whole-cell patch clamp recordings in STIM1/2-KO cells co-expressing Orai1 with either STIM chimera (driven by the robust CMV promoter) demonstrated that S2N-S1C supported a bigger Ca^2+^ release-activated Ca^2+^ (CRAC) current density compared to S1N-S2C (**S4. C, D**). In terms of frequency of Ca^2+^ oscillations, percentage of oscillating cells, plateau cells and non-responding cells, the behavior of these chimeras was solely determined by their C-termini, with tk-S2N-S1C and tk-S1N-S2C supporting similar responses to those of tk-STIM1 and tk-STIM2 respectively (**Fig. 2I-L**). These experiments further demonstrate that STIM1 and STIM2 can independently support Ca^2+^ oscillations and that the C-terminus of STIM1 is the primary driver of plateau responses.

### STIM/Orai isoform interactions shape the diversity of Ca^2+^ signaling events

We recently provided evidence supporting the notion that native CRAC channels are heteromers of Orai isoforms. Without Orai2 and Orai3, physiological Ca^2+^ oscillations typically obtained in response to low- and mid-range agonist concentrations are mostly replaced with Orai1-driven Ca^2+^ plateaus^28^. Here, we performed side by side Förster resonance energy transfer (FRET) experiments with either STIM1-YFP or STIM2-YFP co-expressed with each CFP-Orai isoform in Orai-TKO cells (**Fig. 2M-T**) to gauge the relative interaction between various STIM and Orai isoforms. STIM1 exhibited significant interaction under basal conditions with Orai2 and Orai3 in comparison to Orai1 (**Fig. 2M, N**) and the interaction between STIM1 and Orai isoforms was enhanced after store depletion with thapsigargin to varying degrees (**Fig. 2M-P**). In contrast, STIM2 demonstrated significant basal interactions with Orai1 and Orai2 compared to Orai3 (**Fig. 2Q, R**) and the interaction of STIM2 with each Orai isoform only marginally increased after stimulation with thapsigargin (**Fig. 2Q-T**). Therefore, unlike STIM1 which interacts preferentially at rest with the negative regulators of SOCE, namely Orai2 and Orai3, STIM2 interacts mostly with an isoform that facilitates SOCE (Orai1) and one that dampens SOCE (Orai2). One potential implication from these data is that STIM2 interaction with Orai1 at rest mediates SOCE at low agonist concentrations while interactions of STIM1 with Orai3 dampens SOCE at high agonist concentrations. The interaction of both STIM1 and STIM2 with Orai2 likely fine-tunes SOCE across the full spectrum of physiological agonist concentrations.

To gain insights into how individual STIM proteins communicate with each Orai channel isoform, we used CRISPR/Cas9 gene knockout to generate HEK293 cells expressing combinations of one native Orai isoform and one native STIM isoform. Therefore, we generated six different combinations consisting of the double Orai knockout (Orai2,3-KO, Orai1,2-KO and Orai1,3-KO), in which we knocked out either STIM1 or STIM2 (**S4. B**).

STIM2 knockout in Orai2,3-KO cells caused a slight reduction in SOCE, while STIM1 knockout caused a near complete loss of SOCE (**Fig. 3A, B**). In agreement with previous data from our lab^28^, we showed that under conditions of stimulation with low concentrations of CCh (10 µM), relieving the negative regulatory effects of Orai2 and Orai3 on native Orai1 results in enhanced percentage (∼70%) of Orai2,3-KO cells responding with Ca^2+^ plateaus (**Fig. 3C, Q-S;** compare to 16% of WT-HEK293 cells in **Fig. 1K**). Importantly, in the absence of STIM1, 100% of Orai2,3-STIM1-KO cells responded with Ca^2+^ oscillations (**Fig. 3D, Q-S**). Interestingly, Orai2,3-STIM2-KO cells presented an intermediary phenotype with ∼40% of cells responding with plateaus and ∼60% of cells responding with Ca^2+^ oscillations (**Fig. 3E, Q-S**). Within oscillating cells, deletion of either STIM1 or STIM2 did not affect the frequency of Ca^2+^ oscillations per cell (**Fig. 3P**).

**Fig. 3.**
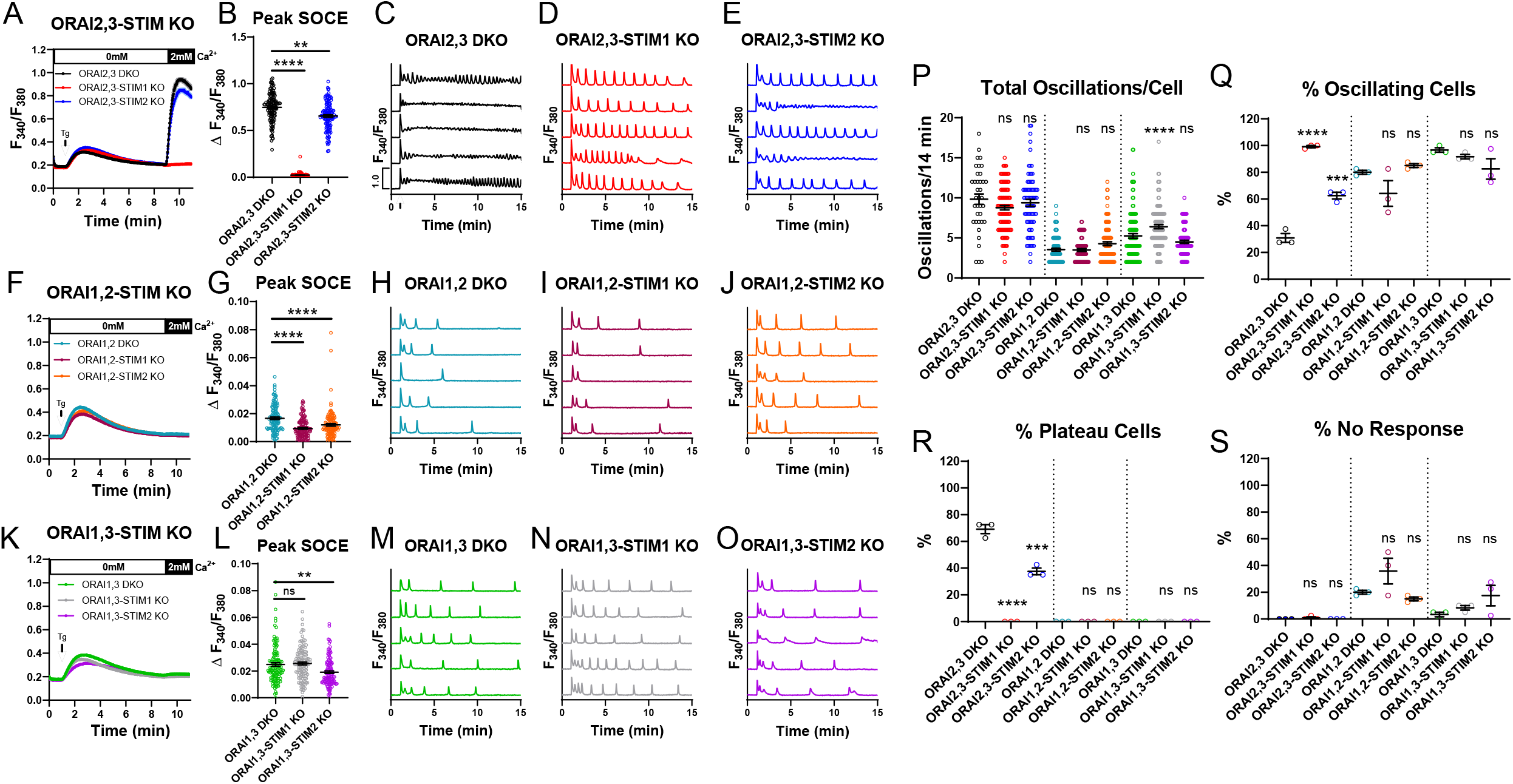
Coordinated STIM/Orai interactions enhance the diversity of Ca^2+^ signaling events. (A) SOCE measurement upon store depletion with 2µM thapsigargin in 0mM Ca^2+^ followed by re-addition of 2mM Ca^2+^ to the bath in Orai2,3-STIM KO cells. (B) Quantification of maximal SOCE for conditions in (A). From left to right *n* = 120, 120, and 120 individual cells. (C-E) Representative Ca^2+^ traces from 5 cells/condition upon stimulation with 10 µM carbachol (CCh) at 1 minute (indicated by arrow in C) in the presence of 2mM Ca^2+^. (F) SOCE measurements of Orai1,2-STIM KO cells. (G) Quantification of maximal SOCE for conditions in (F). From left to right *n* = 120, 120, and 120 individual cells. (H-J) Representative Ca^2+^ traces of Orai1,2-STIM KO cells performed as in (C-E). (K) SOCE measurements of Orai1,3-STIM KO cells. (L) Quantification of maximal SOCE for conditions in (K). From left to right *n* = 120, 121, and 120 individual cells. (M-O) Representative Ca^2+^ oscillation in Orai1,3-STIM KO cells performed as in (C-E). (P) Quantification of total oscillations in 14 minutes for (C-E, H-J, M-O). From left to right *n* = 37, 119, 75, 96, 76, 102, 116, 110, and 99 individual cells. (Q) Quantification of % oscillating cells, (R) % plateau cells, and (S) % non-responding cells for conditions in (C-E, H-J, M-O). For (Q-S) *n* = 3 independent experiments for all conditions. Scatter plots of peak SOCE (B, G, L) and oscillation frequency (P) are presented as mean ± SEM and analyzed with the Kruskal-Wallis one-way ANOVA with multiple comparisons to the respective parental cell line (**p<0.01; ****p<0.0001; ns, not significant). Scatter plots in (Q-S) are presented as mean ± SEM and analyzed with one-way ANOVA with multiple comparisons to the respective parental cell line (***p<0.001; ****p<0.0001; ns, not significant).

Orai1,2-KO cells have an almost complete abrogation of SOCE activated by thapsigargin (**Fig. 3F**). Yet, knockout of either STIM1 or STIM2 in these Orai1,2-KO cells demonstrated an additional very small reduction in SOCE, which only became evident after statistically analyzing many cells (**Fig. 3F, G**). Consistent with our findings that Ca^2+^ plateaus require Orai1^28^, Orai1,2-KO cells lacking either STIM isoform or neither responded strictly with Ca^2+^ oscillations upon stimulation with 10 µM CCh (**Fig. 3H-J, Q-S**). Orai1,2-STIM1-KO cells showed increased percentage of non-responding cells compared to the parental Orai1,2-KO cells, although this increase was not statistically significant (**Fig. 3S**). While Orai1,2-KO and Orai1,2-STIM1-KO cells displayed a similar oscillation frequency (∼3.5 oscillations/14 min), oscillations in Orai1,2-STIM2-KO cells were slightly enhanced (∼4.2 oscillations/14 min) although this did not reach statistical significance (**Fig. 3P**), consistent with preferential STIM1/Orai3 interactions.

Although Orai1,3-KO cells showed an almost complete abrogation of SOCE activated by thapsigargin, the miniscule residual SOCE remaining in these cells was significantly bigger than in Orai1,2-KO cells (**Fig. 3K, L**), consistent with Orai2 displaying higher channel activity than Orai3^28^. STIM2 knockout in Orai1,3-KO cells caused an appreciable decrease of this miniscule SOCE while STIM1 knockout did not (**Fig. 3L**). Orai1,3-KO cells lacking either STIM isoform or neither responded strictly with Ca^2+^ oscillations upon 10 µM CCh stimulation (**Fig. 3M-O, Q-S**). Orai1,3-STIM2-KO cells showed an increased in percentage of non-responding cells compared to the parental Orai1,3-KO cells, although this increase was not statistically significant (**Fig. 3S**). The oscillatory frequency of Orai1,3-KO cells was significantly higher than Orai1,2-KO cells (**Fig. 3P**). STIM1 knockout in Orai1,3-KO cells significantly enhanced the oscillatory frequency of these cells (from ∼5.25 to ∼6.4 oscillations/14 min) while STIM2 knockout caused a small decrease (to ∼4.5 oscillations/14 min) that was not statistically significant (**Fig. 3P**), consistent with preferential STIM2/Orai2 interactions.

### Unactivated STIM inhibit IP_3_R-mediated Ca^2+^ release

The rapid oscillatory response of STIM1/2-KO cells in response to 10 µM CCh (**Fig. 1G, P**) was not observed in Orai-TKO cells (**Fig. 1H**), arguing that this behavior is independent of the loss of SOCE. The frequency of Ca^2+^ oscillations in STIM1/2-KO cells was unchanged in the presence of 5 µM Gd^3+^ (**S2. A-E**), which blocks all Orai channel isoforms^28,41^, ruling out contributions from STIM-independent Ca^2+^ influx through Orai channels. Blockade of Ca^2+^ entry and Ca^2+^ extrusion across the plasma membrane through the use of 1 mM Gd^3+^ in the bath solution, so-called lanthanide insulation^35^, did not alter the Ca^2+^ oscillatory frequencies of WT-HEK293, STIM1-KO, and STIM2-KO cells in response to 10 µM CCh (**Fig. 4A-C, E;** compare **Fig. 4E** to **Fig. 1I**). Importantly, although the Ca^2+^ oscillations of STIM1/2-KO cells were sustained as a result of the lanthanide insulation, their frequency remained rapid (**Fig. 4D, E**), suggesting that the rapid Ca^2+^ oscillations of STIM1/2-KO cells is independent of both Ca^2+^ entry and Ca^2+^ extrusion across the plasma membrane and is likely the result of enhanced ER Ca^2+^ release. Of note, while 10 µM CCh stimulation of STIM1/2-KO cells failed to produce Ca^2+^ plateaus (**Fig. 1K**), when the lanthanide insulation protocol was used ∼ 40% of STIM1/2-KO cells responded with a Ca^2+^ plateau (**S5. B**), suggesting that the combination of enhanced ER Ca^2+^ release in STIM1/2-KO cells and abrogation of Ca^2+^ extrusion increase cytosolic Ca^2+^ enough to produce Ca^2+^ plateau responses.

We performed direct ER Ca^2+^ measurements using the genetically encoded R-CEPIA1*er*. WT-HEK293 cells and their STIM-KO counterparts were stimulated with a supramaximal concentration of CCh (>100 µM) in 0 mM external Ca^2+^ to deplete the ER Ca^2+^ stores, followed by restoration of 2 mM external Ca^2+^ without CCh to allow ER refilling (**Fig. 4F**). Interestingly, both STIM1-KO and STIM2-KO cells displayed a slight increase in the rate of ER Ca^2+^ depletion compared to WT-HEK293 that was not statistically significant. However, we observed a statistically significant increase in the rate of ER Ca^2+^ depletion in STIM1/2-KO cells (**Fig. 4G**). Upon restoration of 2 mM Ca^2+^, the rate of ER Ca^2+^ refilling was significantly reduced in STIM1-KO cells while only marginally reduced in STIM2-KO cells (**Fig. 4H**). As expected, acute ER Ca^2+^ refilling was essentially abrogated in STIM1/2-KO cells (**Fig. 4F, H**). While these data suggest that acute ER refilling at the time scale of minutes is STIM-dependent in agreement with previous work^42^, STIM1/2-KO cells eventually refill their ER Ca^2+^ as documented by the state of their stores at the beginning of recordings. This STIM-independent refilling likely occurs through alternative, and perhaps slower, Ca^2+^ entry routes.

We then utilized two variants of STIM1 and STIM2 that are incapable of activating Orai and mediating SOCE. The first is STIM1 with a point mutation (STIM1-F394H) within the C-terminal STIM-Orai activating region (SOAR)^31^. The second is a naturally occurring STIM2 splice variant called STIM2.1, which does not activate Orai channels^43-45^. STIM1-F394H tagged with CFP and STIM2.1 tagged with YFP were expressed individually and together in STIM1/2-KO cells and ER Ca^2+^ depletion and replenishment were analyzed using the same protocol utilized in **Fig. 4F**. Interestingly, expression of either STIM1-F394H or STIM2.1 reduced the rate of ER Ca^2+^ depletion compared to STIM1/2-KO cells, and this effect was further enhanced with co-expression of both STIM variants (**Fig. 4I, J**). As expected, expression of these STIM variants which are unable to engage and activate Orai proteins, individually or together failed to support acute ER refilling (**Fig. 4I**). In response to 10 µM CCh, STIM1/2-KO cells rapidly oscillate and then cease this activity by ∼5 minutes. Strikingly, rescue with either STIM1-F394H or STIM2.1 prolonged the time of rundown by reducing the frequency of oscillations while increasing their total number (**Fig. 4K-N**). This effect was further enhanced when STIM1-F394H and STIM2.1 were co-expressed in STIM1/2-KO cells (**Fig. 4K, O**). Obviously, because these cells lack both functional Orai-activating STIM1 and STIM2, Ca^2+^ oscillations eventually cease. Rescue with either STIM variant, or both, did not substantially alter the percentages of non-responders, plateau, or total number of oscillating cells (**S5. J-L**). Under conditions where both Ca^2+^ entry and Ca^2+^ extrusion were blocked by lanthanide insulation, co-expression of STIM1-F394H and STIM2.1 in STIM1/2-KO cells caused a reduction in the frequency of oscillations and a decrease in the percentage of cells displaying plateaus (**S5. D-I**), strongly arguing that STIM1/2 N-termini negatively regulate Ca^2+^ release from the ER.

We mutated key residues within the EF-hand of STIM1-F394H (D76A/D78A) and STIM2.1 (D80A/D82A) to abolish their Ca^2+^ binding capacity and mimic STIM activation^12,29^. The expression of these EF-hand mutants in STIM1/2-KO cells either individually or together failed to alter the rate of ER Ca^2+^ depletion in response to supramaximal concentration of CCh (**Fig. 4P, Q**) and failed to alter the oscillatory profile in response to 10 µM CCh (**Fig. 4R-V**), suggesting that unactivated STIMs inhibit Ca^2+^ release. Rescue with either STIM1-F394H (D76A/D78A) and STIM2.1 (D80A/D82A) EF-hand mutants or both did not substantially alter the percentage of non-responders, plateau, or oscillating cells (**S5. M-O**). STIM EF-hand domains are not simply regulating Ca^2+^ release through buffering of ER Ca^2+^. Indeed, ER Ca^2+^ measurements after passive store depletion with the reversible SERCA pump blocker cyclopiazonic acid (CPA, 25 µM) showed that the rate of ER Ca^2+^ depletion by CPA was identical among WT-HEK293, STIM1-KO, STIM2-KO and STIM1/2-KO cells (**S6. A, B**).

We measured the elementary Ca^2+^ signals generated by IP_3_Rs (called Ca^2+^ puffs) in WT-HEK293 cells and STIM1/2-KO cells using TIRF microscopy. Cells were loaded with caged IP_3_ (ci-IP_3_/PM) and Ca^2+^ puffs were recorded after photolysis of ci-IP_3_ for 60 seconds at a rate of 166 frames/second. Ca^2+^ puffs were detected following photolysis of ci-IP_3_ in both WT-HEK293 and STIM1/2-KO cells (**Fig. 5A, B**). The number of puffs and the number of puff sites were significantly higher in STIM1/2-KO cells compared to WT-HEK293 cells (**Fig. 5C,D; Supplementary movie 1** and **2** for WT-HEK293 and STIM1/2-KO cells respectively). Further, Ca^2+^ signals globalized in ∼33 % of STIM1/2-KO cells compared to ∼6 % of WT-HEK293 cells within 60 seconds (**Fig. 5E**), consistent with the increased sensitivity of IP_3_R in STIM1/2-KO cells. The mean rise (r) and fall (f) time of the Ca^2+^ puffs were similar between WT-HEK293 and STIM1/2-KO cells (**Fig. 5F**), indicating that the fundamental biophysical properties of the IP_3_R clusters in each cell type were unaltered. The amplitude distribution of puffs did not differ between WT-HEK293 and STIM1/2-KO cells with most puffs ranging from 0.3 to 0.9 peak amplitude (**Fig. 5G**). Similar results were obtained when Ca^2+^ puffs were evoked by 1 μM CCh (**S7**). The number of puffs and puff sites were significantly greater in STIM1/2-KO as compared to WT-HEK293 cells. While Ca^2+^ signals globalized rarely within 90 s in WT cells (∼9%), ∼45% of STIM1/2-KO cells globalized within this period. The amplitude distribution and mean rise and fall time of the Ca^2+^ puffs were similar between WT-HEK293 and STIM1/2-KO cells (**S7**).

**Fig. 5.**
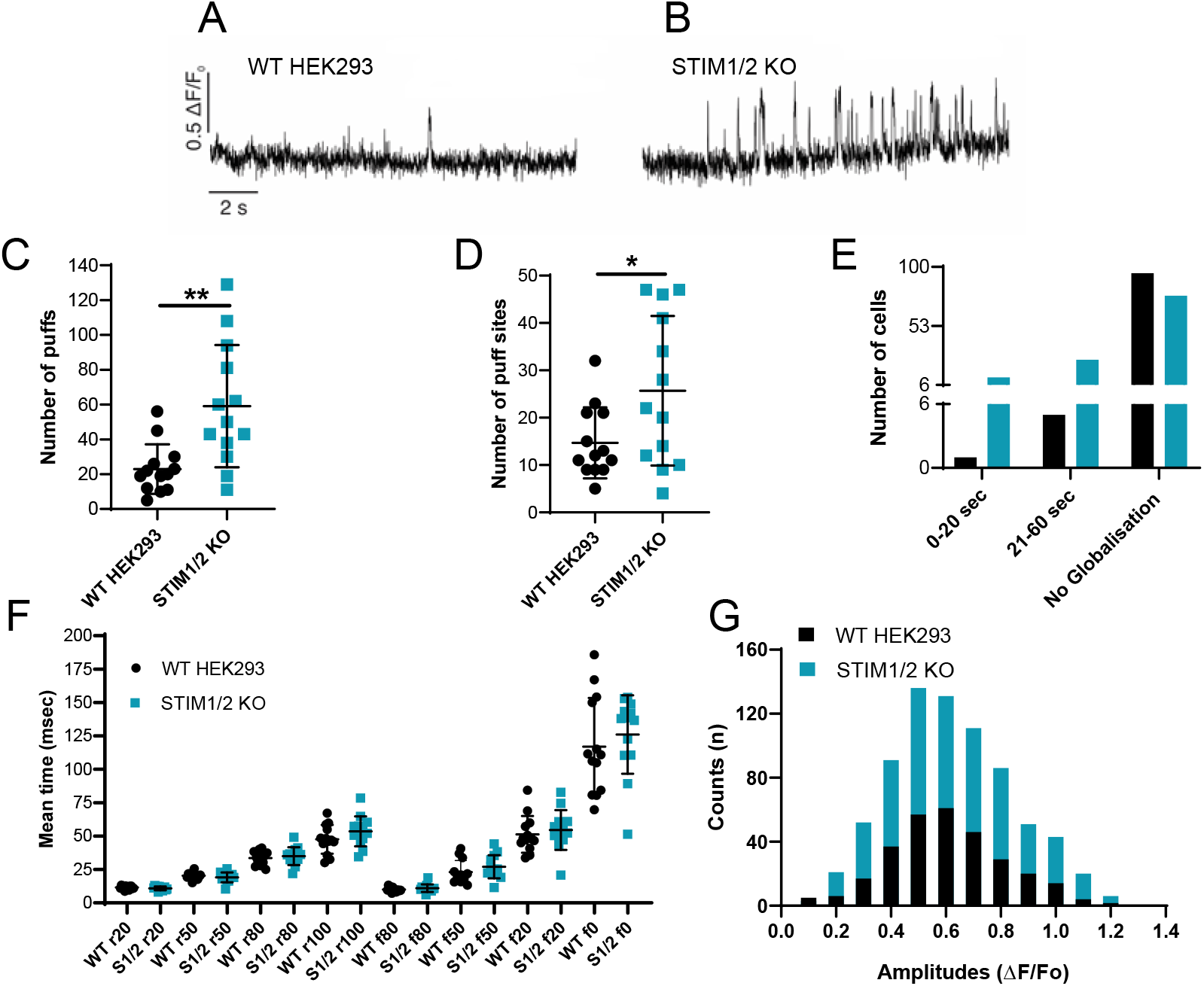
STIM1/STIM2 regulate IP_3_R-mediated Ca^2+^ puffs in response to IP_3_ uncaging. Ca^2+^ puffs were evoked by photolysis of ci-IP_3_ in WT-HEK293 and STIM1/2-KO cells. (A, B) Representative traces showing the Cal-520 fluorescence ratio measurements (ΔF/F_0_) from the center of a single puff site (region of interest 1.3×1.3 µm) evoked following photolysis of ci-IP_3_ in WT-HEK293 (n= 12 independent experiments with 101 cells) and STIM1/2-KO (n= 10 independent experiments with 115 cells), respectively. Photolysis occurred at 3 s, followed by recording for 20 seconds. (C, D) 13 cells from each condition were randomly selected to quantify the number of puffs and puff sites. (C) The number of puffs were significantly higher in STIM1/2-KO (n= 768 in 13 cells) when compared to HEK293 cells (n= 298 in 13 cells). Similarly, (D) the number of puff sites were significantly higher in STIM1/2-KO (n= 334 in 13 cells) when compared to HEK293 cells (n= 191 in 13 cells). (E) From a total of 101 HEK293 cells and 115 STIM1/2-KO cells, bar graph showing the proportions of WT-HEK293 and STIM1/2-KO cells in which the calcium signals globalize within 0-20 seconds or 21-60 seconds following photolysis of ci-IP_3_. Also, shown are the number of cells in which the calcium signals did not globalize. (F) Mean rise and decay times for the fluorescence of Ca^2+^ puffs evoked by photolysis of ci-IP_3_ when it increases or decreases to 20%, 50%, 80%, 100% from 13 cells each of HEK293 and STIM1/2-KO are plotted. (G) Histogram representing the amplitudes distribution of the Ca^2+^ puffs following photolysis of ci-IP_3_ in WT-HEK293 and STIM1/2-KO cells. Data presented as mean ± SEM. Statistical significance was determined by student’s t-test (unpaired, two-tailed). *p<0.05, **p<0.01.

### Mathematical modeling supports STIM1/2, Orai1/2/3 and IP_3_Rs shape receptor-evoked Ca^2+^ signaling

We constructed a simplified model of how SOCE is controlled by the interaction of STIMs (defined as STIMi where i=1,2) with Orai dimers (defined as Oraij,k where j,k = 1,2,3). The Orai model is taken from our recent study^28^, to which we added a simplified model of STIM1 and STIM2 interactions with Oraij,k. This model assumes that both STIMs are activated by a decrease in ER Ca^2+^ and that each STIM can bind to each of the Orai dimer combinations, but with different equilibrium constants. For WT-HEK293, STIM1-KO, STIM2-KO and STIM1/2-KO cells, the model qualitatively reproduces the range of Ca^2+^ oscillations and plateaus in response to increasing concentrations of agonist stimulation (represented by “p”; **Fig. 6A-D**; Solid graphs). Specifically, the model predicts that both STIM1-KO and STIM2-KO cells can support Ca^2+^ oscillations, but that STIM1 is strictly required for Ca^2+^ plateaus (**Fig. 6, A-C**; Solid graphs). The model reproduces the high sensitivity to agonist and the faster oscillatory frequency of STIM1/2-KO cells (**Fig. 6D**; Solid graphs), in support of the regulation of IP_3_-mediated Ca^2+^ release from the ER by unactivated STIM proteins. The model reproduces the experimental data obtained with the Gd^3+^ insulation protocol (i.e. closed cell system; **S8. A-D**). STIM1/2-KO cells readily respond by Ca^2+^ plateaus when Ca^2+^ extrusion is abrogated in the closed cell condition (**S8. D**). The model qualitatively reproduces the experimental data on ER Ca^2+^ measurements (**Fig. 6E, F**; depicting normalized and absolute ER Ca^2+^ levels, respectively). Under these simulations, all the STIM-KO cells show faster ER Ca^2+^ depletion, while ER refilling rates of STIM1-KO and, more so, that of STIM1/2-KO cells are drastically slowed (**Fig. 6E, F**).

**Fig. 6.**
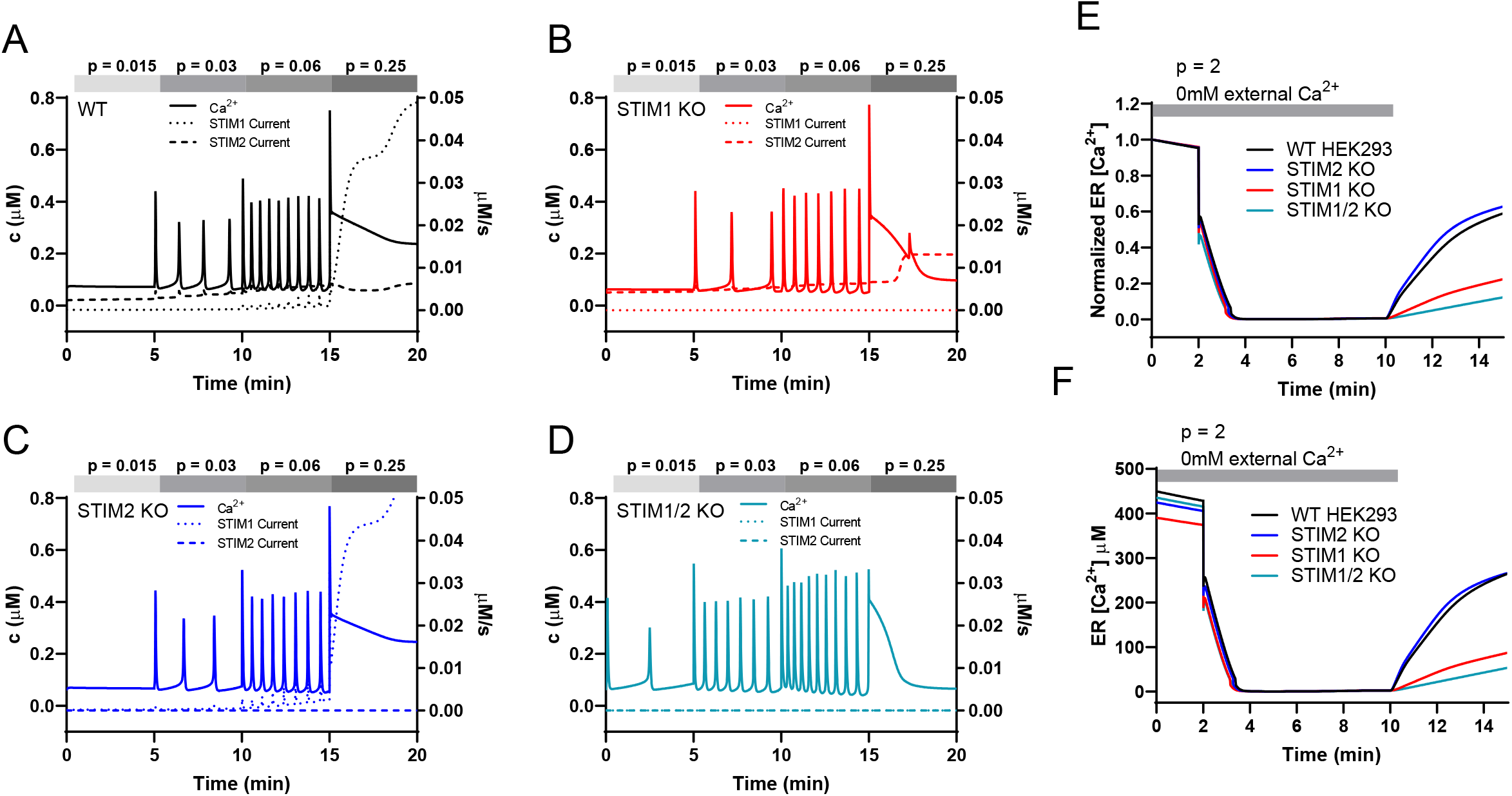
Mathematical modeling supports coordinated STIM1/STIM2 functions over the diverse range of Ca^2+^ signaling events. Responses to four different agonist concentrations (*p* = 0.015, *p* = 0.03, *p* = 0.06, *p* = 0.25) applied sequentially in that order, each for 5 min. In each panel the solid line is the Ca^2+^ concentration, and is plotted against the left axis, while the dotted and dashed lines are the STIM currents, which are plotted against the right axis. A-D; Ca^2+^ responses under physiological extracellular Ca^2+^ conditions (open cell), with all three Orai isoforms present. The STIM1/2-KO cell is the most effective oscillator (as seen in **Fig. 1**), which results from the model assumption that the IP_3_R is inhibited by unactivated STIM. In the absence of STIM1 (B and D) the cell is unable to maintain raised Ca^2+^ plateaus, even at high agonist concentrations.

The experimental data show that at the beginning of recordings the ER Ca^2+^ stores of STIM1/2-KO cells are essentially full (**Fig. 4, F, I, P**; **Fig. 5, E, F**; see also^42^), suggesting the presence of an alternative STIM-independent pathway that can refill ER Ca^2+^ in a matter of hours or days. The nature of this STIM-independent pathway is not clear currently. As such, the model predicts that under basal conditions (i.e., without agonist), only 28% of the Ca^2+^ influx into the cell is through STIM/Orai channels; 7% (of this 28%) of STIM/Orai-mediated current is mediated by STIM1, with the remaining 93% mediated by STIM2 (**Fig. 6**; dotted graphs). For low agonist concentrations this proportion is maintained. At higher agonist concentrations, 92% of the Ca^2+^ influx into the cell is mediated by STIM/Orai channels, and of that, 93% is mediated by STIM1 and 7% by STIM2. The model supports the experimental data showing that both STIM1 and STIM2 can support oscillations and are both involved across the full range of agonist concentrations. However, in the absence of STIM1, even high agonist concentrations do not lead to sustained plateaus.

Further, the model supports the experimental data (**Fig. 3**) obtained with Orai double knockout/STIM single knockout cells. The simulations (depicted in **S9**) show the responses of cells expressing combinations of one native Orai and one native STIM isoform to stimulation with thapsigargin, first in 0 mM Ca^2+^, then in Ca^2+^-containing external solution. Model simulations of the application of increasing concentrations of agonist to these cells in Ca^2+^-containing external solution qualitatively reproduce the experimental data showing that native STIM1 and Orai1 are essential for mediating Ca^2+^ plateaus in response to high agonist concentrations with STIM2, Orai2 and Orai3 regulating SOCE across the full range of agonist concentrations, but have negligible contributions to plateaus on their own (**S9**). Consistent with these results, model simulations show that in WT-HEK293 cells, at all agonist concentrations, STIM1 interaction with the Orai1,1 homodimer mediates the great majority of SOCE activity (**S10**). Finally, the model reproduces the dynamic change of oscillation properties in response to agonist application in WT-HEK293 cells, as the oscillations initially start faster but gradually change to slower oscillations on a lower baseline (see e.g. **S1. H**), a known feature of Ca^2+^ oscillation models^46,47^.

### Coordinated STIM1/STIM2 functions tailor receptor-activation to NFAT induction

SOCE is a critical regulator of NFAT isoform nuclear translocation^14,18,48-51^. The nuclear translocation of NFAT1 is relatively slow and requires robust SOCE activity, while NFAT4 nuclear translocation is more rapid and requires relatively low SOCE activity^28,48,52-54^. We expressed GFP-tagged NFAT1 and NFAT4 reporter constructs in WT-HEK293, STIM1-KO, STIM2-KO and STIM1/2-KO cells. These four cell types were then stimulated with either low (10 µM) or high (300 µM) concentration of CCh and NFAT isoform nuclear translocation was monitored. In WT-HEK293 cells, stimulation with 10 µM CCh caused a slow and gradual accumulation of NFAT1 in the nucleus (**Fig. 7A, E**). NFAT1 nuclear translocation was significantly reduced in STIM1-KO and STIM2-KO cells and essentially abolished in STIM1/2-KO cells (**Fig. 7A, B, E**). When WT-HEK293 cells were stimulated with 300 µM CCh, NFAT1 nuclear translocation was more robust (**Fig. 7C**). While NFAT1 nuclear translocation was completely inhibited in STIM1/2-KO cells, it was only slightly inhibited in STIM2-KO cells and substantially reduced in STIM1-KO cells (**Fig. 7C, D, F**). The nuclear translocation of NFAT4 in WT-HEK293 cells was significantly faster by comparison to that of NFAT1 at both agonist concentrations (**Fig. 7G-L**). NFAT4 nuclear translocation in response to either 10 µM or 300 µM of CCh was significantly reduced in STIM1-KO and STIM2-KO cells and essentially abrogated in STIM1/2-KO cells (**Fig. 7G-L**). These results demonstrate that both STIM1 and STIM2 are required for optimal NFAT nuclear translocation under both low and high agonist stimulus intensities.

**Fig. 7.**
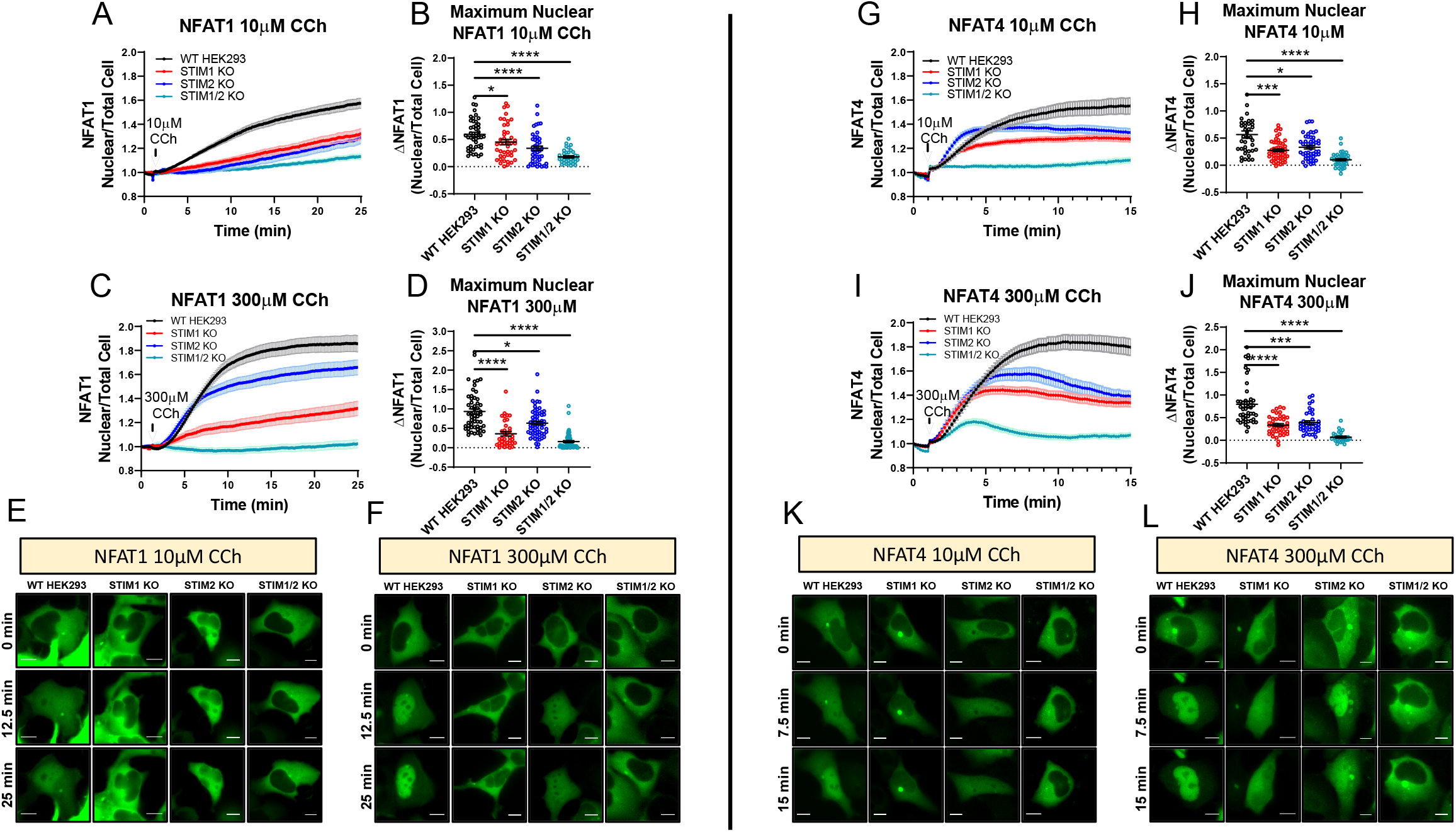
Both STIM1 and STIM2 are required for optimal NFAT1/4 activation. (A) Time-lapse quantification of NFAT1-GFP nuclear translocation in response to 10µM CCh in 2mM Ca^2+^ as determined by the ratio of nuclear/total GFP fluorescence. (B) Quantification of maximal nuclear NFAT1 fluorescence for each condition in (A) at 25 minutes. From left to right *n* = 46, 43, 44, and 36 individual cells. (C) Quantification of NFAT1-GFP nuclear translocation in response to 300µM CCh. (D) Quantification of maximal nuclear NFAT1 fluorescence from conditions in (C) at 25 minutes. From left to right *n* = 55, 36, 62, and 59 individual cells. (E) Representative time-lapse images of NFAT1-GFP expressing cells after 10µM CCh or (F) 300µM CCh stimulation. (G) Same conditions as in (A) but utilizing NFAT4-GFP expressing cells. (H) Quantification of maximal nuclear NFAT4 fluorescence for conditions in (G) at 15 minutes. From left to right n = 39, 53, 49, and 40 individual cells. (I) Quantification of NFAT4-GFP nuclear translocation in response to 300µM CCh. (J) Quantification of maximal nuclear NFAT4 fluorescence from conditions in (I) at 15 minutes. From left to right *n* = 46, 40, 36, and 29 individual cells. (K) Representative time-lapse images of NFAT4-GFP expressing cells after 10µM CCh or (L) 300µM CCh. Scale bar = 10 µm. Scatter plots of maximal nuclear NFAT1 (B, D) and NFAT4 (H, J) are presented as mean ± SEM and analyzed with the Kruskal-Wallis one-way ANOVA with multiple comparisons to WT-HEK293 (*p<0.05; ***p<0.001; ****p<0.0001; ns, not significant).

## Discussion

It is clearly established that SOCE in response to agonist stimulation requires the interaction between STIM1 and Orai1 and the molecular mechanisms underlying their coupling has been intensively investigated^10,55-58^. It is also known that STIM2 is a weaker SOCE activator and its recruitment is triggered under basal conditions, when ER Ca^2+^ store depletion is modest, to maintain homeostatic ER Ca^2+^ levels^29^. Recently, we showed that all three Orai isoforms oligomerize to form native CRAC channels, enhancing the diversity of Ca^2+^ signaling events in response to the full range of stimulus intensities^28,59^. Here, we addressed the choreography of interactions between each native STIM1/2 and Orai1/2/3. Multiple caveats have hindered the understanding of the contributions of each native STIM/Orai to mammalian Ca^2+^ signaling. First, ectopic expression has been performed on a wildtype background, such that the interference of endogenous SOCE is unknown. Ectopic expression typically produces protein levels that are higher than endogenous levels^48^, altering stoichiometry and channel activity. In most of these overexpression studies, only STIM1 and Orai1 were considered. Second, measurements of native SOCE activity have been mostly performed with protocols that maximal store depletion^60^. Under physiological conditions, stimulation of cells with relatively low or moderate agonist concentrations induces concomitant IP_3_R-mediated Ca^2+^ oscillations and ER Ca^2+^ replenishment from the extracellular space through SOCE, resulting in only minor and transient fluctuations of ER Ca^2+^ content. Third, prior reports specifically investigating the role of native STIM1 and STIM2 in Ca^2+^ oscillations triggered by relatively low agonist concentrations have relied on siRNA knockdown^33-36^. These experiments produced conflicting results regarding the relative contributions of STIM1 versus STIM2. Variations in knockdown efficiencies between studies are likely confounding factors.

To overcome these limitations, we investigated Ca^2+^ signaling in multiple clones of HEK293 cells devoid of STIM1 or STIM2 proteins individually and in combination. In response to stimulation with low concentrations of agonist, both STIM1-KO and STIM2-KO cells maintain oscillation frequencies like WT-HEK293 cells. However, loss of STIM1 completely abolished Ca^2+^ plateaus. STIM2-KO cells exhibit plateaus at lower concentrations of agonist in comparison to WT-HEK293, suggesting that STIM2 negatively regulates SOCE activity and is required for enhancing the repertoire of Ca^2+^ signaling events. Ca^2+^ plateaus are driven exclusively by the C-terminus of STIM1. Alterations in Ca^2+^ oscillation profiles are predicted to have important downstream functional consequences, particularly in the activation of the Ca^2+^-dependent transcription factors, NFAT^18,48,61^. By investigating the activation of two NFAT isoforms with unique nuclear translocation requirements^52,53^, we demonstrate that loss of either STIM1 or STIM2 impairs NFAT1 and NFAT4 translocation in response to both low (10 µM CCh) and high (300 µM CCh) agonist concentrations, suggesting that the presence of both native STIM1 and STIM2 is always required for optimal activation of both NFAT1 and NFAT4 at all agonist concentrations. The combined functions of STIM1 and STIM2 are required to tailor the Ca^2+^ signal to the strength of agonist stimulation and NFAT transcriptional activity. STIM1 plays the major role for NFAT1 nuclear translocation in response to high concentrations of agonist. This is consistent with the known properties of NFAT1, which requires robust Ca^2+^ concentrations for activation and with STIM1 function described herein as the exclusive driver of Ca^2+^ plateaus. We recently revealed that coordination between endogenous STIM1 and STIM2 plays a critical role in mediating SOCE in the colorectal cancer cell line HCT116^40^. Further, recent studies presented evidence that STIM2 can serve as an enhancer of receptor-activated Ca^2+^ signaling by recruiting STIM1 to Orai1 channels under conditions of modest store depletion^33,37,38^, in agreement with our findings. Whether STIM1 and STIM2 function as homodimers or heterodimers is an important question that requires further investigations. Native CRAC channels are likely heterohexamers of all three Orais and STIM1 displays stronger basal interactions with Orai2 and Orai3 compared to Orai1^28,62^. Interestingly, STIM2 shows stronger basal interaction with Orai1 and Orai2 compared to Orai3. Our combinations of single STIM-KO/double Orai-KO and modeling defined the role of native STIM/Orai duos in shaping Ca^2+^ signaling events. Ca^2+^ plateaus strictly require the STIM1/Orai1 duo, which also mediate the diversity of Ca^2+^ oscillation frequencies in combination with the other five STIM/Orai duos.

Although STIM1/2-KO cells completely lack SOCE, their oscillation profile does not phenocopy that of Orai-TKO cells. STIM1/2-KO cells are more sensitive to IP_3_-mediated Ca^2+^ release and we show that STIM1/2 negatively regulate IP_3_R channel activity. While the mechanisms of this IP_3_R regulation by STIMs require further experimentation, our mutagenesis, ER Ca^2+^ measurements and modeling data suggest it is the unactivated STIMs that inhibit IP_3_R channels, likely setting the threshold of activation by IP_3_. As STIM proteins unfold and become increasingly activated (a state mimicked by EF-hand mutants), the inhibition of IP_3_Rs is lifted with concomitant and equivalent CRAC channel activation, thus precisely coordinating the levels of Ca^2+^ release and Ca^2+^ entry. Taken together, our work clarifies the complex and dynamic choreography of native interactions between IP3Rs, two STIM and three Orai proteins and how this set of proteins provides the mammalian cell with an effective toolkit to enhance the diversity of Ca^2+^ signaling events and subsequent transcriptional control in response to the full spectrum of agonist intensities.

## Supporting information

WT-HEK293_Ca2+ puffs

STIM1-2-KO-Ca2+ puffs

List of reagents

Raw data

Stastics

## Data Availability

All materials and experimental protocols; all raw data and raw unprocessed gels and Westerns; the source and reference number of reagents, recombinant DNA and animals necessary for replication are included in an online appendix. The exact “p values” for all statistical comparisons are also included in an online appendix.

## Acknowledgments

We are grateful to Drs. Jeff Lock and Ian Parker from The University of California, Irvine for their advice and help with the Ca^2+^ puff measurements using TIRF microscopy and with data analysis.

## Author contributions

SME, REY, DIY and MT designed research; SME, REY, XP, VA and LEW performed research; JS conceived and performed mathematical simulations; DIY, NH and DLG contributed new reagents/analytic tools; SME, REY, VA, DIY and MT analyzed data; and SME and MT wrote the paper with input from all authors.

