## Supplementary material for "Omnitemporal choreographies of IP_3_R and all five STIM/Orai underlie the complexity of mammalian Ca^2+^ signaling": Supplemnetal Figures and methods

*By*

Scott M. Emrich[*,1], Ryan E. Yoast[*,1], Ping Xin[1], Vikas Arige[4], Larry E. Wagner[4], Nadine Hempel[2,3], Donald L. Gill[1], James Sneyd[5], David I. Yule[4], and Mohamed Trebak[#,1,3]

### Methods

#### Cell culture and transfection

All HEK293 cell lines were cultured in high glucose (4.5g/L) Dulbecco's modified Eagle's medium (DMEM) supplemented with 10% fetal bovine serum, 1X Antibiotic-Antimycotic (Thermo Scientific), and cultured at 37°C with 5% $CO_2$. For CMV-driven STIM and Orai constructs, HEK293 cells were transfected using an Amaxa Nucleofector II (Amaxa Biosystems) with the Nucleofector Kit V (VVCA-1003) and program Q-001. For TK-driven and NFAT reporter constructs, cells were transfected using Lipofectamine 2000 (Invitrogen) following the manufacturer's protocol. Following each transfection cells were plated on 25mm glass coverslips and incubated for 24 hours before use in experiments.

#### Generation of STIM and Orai CRISPR/Cas9 knockout cell lines

HEK293 Orai and STIM knockout cell lines were generated as previously described [1-3]. Briefly, sequence specific guide RNAs (gRNA) targeting *STIM1*, *STIM2*, *Orai1*, *Orai2*, or *Orai3* were cloned into one of two fluorescent vectors (pSpCas9(BB)-2A-GFP, Addgene #48138; pU6-(BbsI)_CBh-Cas9-T2A-mCherry, Addgene #64324). For genes with commercially available validated antibodies (*STIM1*, *STIM2*, and *Orai1*) a single gRNA was utilized to reduce off target effects. For genes without validated antibodies (*Orai2* and *Orai3)*, two gRNA sequences were used to excise the entire gene. Parental HEK293 cell lines were transfected with each fluorescent CRISPR vector and individual single or double GFP/mCherry positive cells were FACS sorted (BD FACS Aria SORP) into 96-well cell culture plates 24 hours after transfection. Individual knockout clones were validated using western blot analysis, Sanger sequencing, and functional $Ca^{2+}$ imaging experiments. Each gRNA sequence used to generate single and double Orai and STIM knockout cell lines was as follows:

- *Orai1* g1: 5'-GTTGCTCACCGCCTCGATGT-3'

- *Orai2* g1: 5'-ACGACAGGGCCTGTACCGAG-3'
- *Orai2* g3: 5'-CTCATGCGGGGACTCGCTGA-3'
- *Orai3* g2: 5'-GTTCGTGCACCGCGGCTACC-3'
- *Orai3* g4: 5'-CCAGAGACTGCACCGCTACG-3'
- *STIM1* g3: 5'-TGATGAGCTTATCCTCACCA-3'
- *STIM2* g1: 5'-AGATGGTGGAATTGAAGTAG-3'

**DNA Constructs**

All novel tk-driven constructs (tk-STIM1, tk-STIM2, tk-S1N-S2C, tk-S2N-S1C) and STIM1/STIM2 EF-hand mutants were generated with primers compatible with the In-Fusion cloning system (Takara Bio Inc.). Primers used to create the D80A/D82A substitutions in YFP-STIM2.1 were 5'-ACAAATGGCTGATGCCAAAGATGGTGGAATTG-3' and 5'-TGGCATCAGCCATTTGTTTATGTATTGTTTGAAG-3'. Primers used to create the D76A/D78A substitutions in STIM1-F394H-YFP were 5'-ACTGATGGCCGATGCTGCCAATGGTGATGTGG-3' and 5'-CAGCATCGGCCATCAGTTTGTGGATGTTACGG-3'. CMV-CFP-Orai1, CMV-CFP-Orai2, CMV-CFP-Orai3, STIM1-YFP, STIM2-YFP, YFP-STIM2.1, and STIM1-F394H-YFP were from Dr. Don Gill. HA-NFAT1(4-460)-GFP (Addgene #11107) and HA-NFAT4(3-407)-GFP (Addgene #21664) were from Dr. Anjana Rao. pCMV R-CEPIA1er was from Dr. Masamitsu Iino (Addgene #58216).

**Fluorescence Imaging**

HEK293 cells ($1.5x10^5$) were plated onto 25mm glass coverslips and allowed to attach overnight. For cytosolic $Ca^{2+}$ measurements, coverslips were mounted in Attofluor cell chambers (Thermo Scientific) and cells were incubated in complete DMEM containing 2μM Fura-2AM (Molecular Probes) for 30 minutes at 37°C. Following Fura-2 loading, cells were washed 3 times with a HEPES-buffered saline solution (HBSS) containing the following components: 120mM NaCl, 5.4mM KCl, 0.8mM $MgCl_2$, 2mM $CaCl_2$, 20mM Hepes, 10mM D-glucose, and adjusted to pH 7.4 with NaOH. Chambers were mounted on a Leica DMi8 fluorescence microscope and Fura-2 fluorescence was measured every 2 seconds by excitation at 340nm and 380nm using a fast shutter wheel and the emission at 510nm was collected through a 20X fluorescence objective. Fluorescence data was collected from individual cells on a pixel by pixel basis and processed using Leica Application Suite X. All cytosolic $Ca^{2+}$ concentrations are presented as the ratio of $F_{340}/F_{380}$ and plotted as means ± SEM from at least three independent experiments.

For ER $Ca^{2+}$ measurements, HEK293 cell lines were transfected by Nucleofection with 1μg of R-CEPIA1er and imaged 24 hours later on a Leica TCS SP8 confocal microscope with a 40X oil objective. RFP fluorescence was excited with the 552nm laser at 1% output at room temperature. Images were collected every 10-15 seconds and traces were plotted as means ± SEM after normalization to the first timepoint. Prior to the start of each recording, cells were switched from HBSS containing 2mM $CaCl_2$ to a HBSS solution nominally free of $CaCl_2$.

For NFAT nuclear translocation, HEK293 cell lines were transfected with 1μg of NFAT1-GFP or NFAT4-GFP reporter constructs using Lipofectamine 2000. Cells were imaged 24 hours post transfection using a Leica DMi8 fluorescence microscope equipped with a Hamamatsu Flash 4 camera with a 40X oil objective. GFP fluorescence from the whole cell and nucleus was collected every 10 seconds using a 488nm fast filter wheel with emission captured through a corresponding GFP filter cube. NFAT1 and NFAT4 nuclear translocation was calculated using the equation $NFAT\ translocation = (\frac{Nuclear\ F_{510}}{total\ cell\ F_{510}})$ and plotted as a function of time in response to carbachol stimulation.

**Förster resonance energy transfer (FRET) measurements**

Measurement of FRET between STIM1 and STIM2 with each Orai isoform was performed as previously described[3]. Briefly, Orai-TKO cells were transfected by Nucleofection with 2µg STIM1-YFP in combination with 3µg CFP-Orai1, 4µg CFP-Orai2 or 5µg CFP-Orai3 and 2µg STIM2-YFP together with 3µg CFP-Orai1, 4µg CFP-Orai2, or 5µg CFP-Orai3. Imaging was performed on a Leica DMI 6000B inverted automated fluorescence microscope equipped with a 40X oil objective with simultaneous collection of CFP, YFP, and FRET images every 20 seconds. Exposure times for STIM1 and STIM2 FRET were 1000ms (CFP), 250ms (YFP), and 1000mM (FRET). Three channel FRET was calculated with SlideBook 6.0 software (Intelligent Imaging Innovations) using the formula $F_C = I_{DA} - \frac{Fd}{Dd} \text{ x } I_{DD} - \frac{Fa}{Da} \times I_{AA}$ where, $I_{DD}$, $I_{AA}$, and $I_{DA}$ were the respective fluorescent intensities of CFP, YFP, and FRET channels after taking individual CFP and YFP background correction images. Calculation of E-FRET was performed as previously described using the formula $E_{app} = Fc\frac{Fc}{Fc+G\ x\ I_{DD}}$ [4].

**Detection and analysis of $Ca^{2+}$ puffs using TIRF microscopy**

Parental WT-HEK293 cells or STIM1/2-KO cells were grown on 15-mm glass coverslips coated with poly-D-lysine (100 µg/ml) in a 35-mm dish for 2 days. Prior to imaging, the cells were washed three times with imaging buffer (137 mM NaCl, 5.5 mM glucose, 0.56 mM $MgCl_2$, 4.7 mM KCl, 1.26 mM $CaCl_2$, 10 mM HEPES, 1 mM $Na_2HPO_4$ at pH 7.4). The cells were subsequently incubated with Cal520-AM (5 µM; AAT Bioquest #21130) and ci-$IP_3$/PM (1 µM, Tocris #6210) in imaging buffer with 0.01 % BSA in dark at room temperature. After 1 hour incubation, the cells were washed three times with imaging buffer and incubated in imaging buffer containing EGTA-AM (5 µM, Invitrogen #E1219). After 45 minutes incubation, the media was replaced with fresh imaging buffer and incubated for additional 30 minutes at room temperature to allow for de-esterification of loaded reagents[5]

Following loading, the coverslip was mounted on chambers and imaged using an Olympus IX83 inverted total internal reflection fluorescence microscopy (TIRFM) equipped with oil-immersion PLAPO OTIRFM 60x objective lens/1.45 numerical aperture. The cells were illuminated using a 488 nm laser to excite Cal-520 and the emitted fluorescence was collected through a band-pass filter by a Hamamatsu ORCA-Fusion CMOS camera. The angle of the excitation beam was adjusted to achieve TIRF with a penetration depth of ~140 nm. Images were captured from a field of view by directly streaming into RAM. To photorelease $IP_3$, UV light from a 405 nm laser was introduced to uniformly illuminate the field of view. Both the intensity of the UV flash (~200 µW) and the duration (1000 msec) for uncaging $IP_3$ were optimized to prevent spontaneous puff activity in the absence of loaded ci-$IP_3$. TIRF images were captured using 4 X 4 pixel binning (433.333 nm/pixel) from equal field of views for WT-HEK293 and STIM1/2-KO cells at a rate of ~166 frames per second. After visualizing images with the cellSens [Ver.2.3] life science imaging software (Olympus), images were exported as vsi files. Images, 10 seconds before and 60 seconds after flash photolysis of ci-$IP_3$, were captured.

The vsi files were converted to tif files using Fiji and further processed using FLIKA, a Python programming based tool for image processing[6]. From each recording, 500 frames (~3 seconds) before photolysis of ci-$IP_3$ were averaged to obtain a ratio image stack (F/Fo) and standard deviation for each pixel for recording up to 20 seconds following photolysis. The image stack was Gaussian-filtered, and pixel that exceeded a critical value (1.0 for our analysis) were located. The 'Detect-puffs' plug-in was utilized to detect the number of clusters (puff sites), number of events (number of puffs), amplitudes and durations of localized $Ca^{2+}$ signals from individual cells. All the puffs identified automatically by the algorithm were manually confirmed before analysis[7,8]. The results from FLIKA were saved as excel and graphs were plotted using GraphPad Prism 8.

A similar protocol for loading was followed to test the effect of CCh on $Ca^{2+}$ puffs. In this case, the cells were not loaded with ci-$IP_3$. Cells were stimulated by perfusing 1 μM CCh to generate $IP_3$-induced $Ca^{2+}$ release.

**Western blot analysis**

HEK293 cell lines were harvested and lysed for 15 minutes in ice cold RIPA buffer (150 mM NaCl, 1.0% IGEPAL CA-630, 0.5% sodium deoxycholate, 0.1% SDS, 50 mM Tris, pH 8.0; Sigma) containing 1X Halt protease/phosphatase inhibitors (Thermo Scientific). Following lysis, samples were clarified by centrifugation at 15,000xg for 10 minutes at 4°C. Supernatants were collected and protein concentration was determined using the Pierce Rapid Gold BCA Protein Assay Kit (Thermo Scientific). Equal concentrations of protein extract were loaded into 4-12% NuPAGE BisTris gels (Life Technologies) and transferred to PVDF membranes utilizing the Transblot Turbo Transfer System (Bio-Rad). Membranes were blocked for 1 hour at room temperature in Odyssey Blocking Buffer in TBS (LI-COR) and incubated overnight at 4°C with primary antibody. The following antibodies and dilutions were used: STIM1 (1:2000; 4916, Cell Signaling Technology), STIM2 (1:1000; 4917, Cell Signaling Technology), α-tubulin (1:5000; 3873, Cell Signaling Technology), GAPDH (1:5000; MAB374, Sigma), IP3R-1 (1:1000, Yule Lab), IP3R-2 (1:1000, Yule Lab), and IP3R-3 (1:1000; 610313, BD Biosciences). Membranes were washed with TBST and incubated for 1 hour at room temperature with the following secondary antibodies: IRDye 680RD goat anti-mouse (1:10,000 LI-COR) or IRDye 800RD donkey anti-rabbit (1:10,000 LI-COR). Membranes were imaged on an Odyssey CLx Imaging System (LI-COR) and analysis performed in Image Studio Lite version 5.2 (LI-COR) and ImageJ.

**Patch clamp electrophysiology**

For measurement of $I_{crac}$, STIM1/2-KO cells were transfected by Nucleofection with 3.5μg of each YFP-tagged STIM chimera (YFP-S1N-S2C or YFP-S2N-S1C) along with 1.5μg of CFP-Orai1. Cells were seeded onto 30mm glass coverslips and patch clamp recordings were performed 24 hours after transfection. Patch-clamp recordings were performed using an Axopatch 200B along with a Digidata 1440A (Molecular Devices). Pipettes were pulled from borosilicate glass capillaries using a P-1000 Flaming/Brown micropipette puller (Sutter Instrument Company) and a DMF1000 (World Precision Instruments) was used for polishing. Glass electrodes with a resistance of 2 to 4 MΩ were utilized. Recordings were only performed when a >16 GΩ seal was formed and when whole cell series resistance was <8 MΩ. Patch Master (HEKA) was used for data acquisition and analysis. For recording of CRAC currents, cells were maintained at a +30mV holding potential and reverse voltage ramps from +100 to -140mV for 250ms at 2s intervals were performed. After establishing the whole-cell configuration, intracellular stores were depleted using 20mM BAPTA. The following pipette and bath solutions were utilized:

Pipette solution: 145 mM cesium methanesulfonate, 20 mM BAPTA, 10 mM Hepes, 8 mM $MgCl_2$, and 25 µM $IP_3$ (pH to 7.2 with CsOH).

Bath solution: 115 mM Na-methanesulfonate, 10 mM CsCl, 1.2 mM $MgSO_4$, 10 mM Hepes, 20 mM $CaCl_2$, and 10 mM glucose (pH 7.4 with NaOH).

**Mathematical modelling**

The majority of the model parameters remain unchanged from [3]. The only additional parameters are the equilibrium constants of the STIMi-Oraij,k interactions and the unitary current through a single STIMi-Oraij,k complex.

The following model assumptions are important for obtaining the correct qualitative behaviour. There is flexibility in the precise numbers used, but the qualitative trends are critical.

1. There is a background STIM-independent influx of $Ca^{2+}$ into the cell, which we call a leak. Knockout of STIM changes the size of this leak, which in turn sets the resting ER $Ca^{2+}$ concentration. However, changes in the leak are not always sufficient to maintain 100% of the basal ER $Ca^{2+}$ concentration when STIM1 or STIM2 are knocked out. STIM2-KO, STIM1-KO and STIM1,2-KO cells have resting ER $Ca^{2+}$ of, respectively, 94%, 86% and 97% of basal. We propose no mechanism for how the leak changes. However, the counterintuitive assumption that STIM1,2-DKO cells have a higher resting ER $Ca^{2+}$ than the STIM1-KO cells is consistent with the findings of Zheng et al [9].
2. Unactivated STIM1 and STIM2 inhibit $IP_3R$. This is modelled by a phenomenological function (see Modeling Methods below) that increases $IP_3R$ activity as the amount of unactivated STIM1 and/or STIM2 decreases, as a result of activation or knockout of STIM. This is a positive feedback loop (agonist stimulation decreases ER $Ca^{2+}$, which activates STIM, which decreases the amount of unactivated STIM, which removes inhibition from the $IP_3R$, which lowers ER $Ca^{2+}$ further), but plays no significant role in the generation of single oscillations. Its main function is to increase oscillation frequency in a STIM1,2 DKO cell that has been closed by 1mM $Gd^{3+}$ (see **Fig. 4 D**).
3. Binding ratios to the Orai homodimers (for example, the ratio of STIM1/Orai1,1 to STIM1/Orai2,2 binding, and so on) are determined from the FRET data in **Fig. 2 M, Q**; see details below in Modeling methods.
4. There is three times more STIM1 than STIM2.

5. In most cases, the unitary current through a STIM2-Oraij,k complex is about 1/5 of the current through the analogous STIM1-Oraij,k complex. The two exceptions are the unitary currents through STIM2-Orai2,2 and STIM2-Orai3,3 which are ten times the current through STIM1-Orai2,2 and STIM1-Orai3,3 respectively.

**Modeling Methods:** The models of Sneyd et al and Yoast et al[3,10] are combined with a simple STIM model. Let $S_1$ denote the concentration of STIM1 and let $S_1^*$ denote the concentration of activated STIM1, with analogous notation for STIM2. Assuming that $S_1$ is activated in the reaction

$$S_1^* \underset{k_{-1}}{\overset{k_1 c_e^2}{\rightleftharpoons}} S_1$$

it follows that, at steady state,

$$\frac{S_1^*}{S_1^T} = \frac{K_{e1}^2}{K_{e1}^2 + c_e^2} = \phi_1(c_e),$$

where $S_1^T$ is the maximal concentration of STIM1, and where $K_{e1}^2 = \frac{k_{-1}}{k_1}$. Similarly,

$$\frac{S_2^*}{S_2^T} = \frac{K_{e2}^2}{K_{e2}^2 + c_e^2} = \phi_2(c_e).$$

The STIM bind to the various Orai dimers as follows,

$$S_1^* + O_{ij} \rightleftharpoons P_{ij},$$
$$S_2^* + O_{ij} \rightleftharpoons Q_{ij},$$

and thus, at steady state,

$$P_{ij} = p_{ij}\phi_1(c_e)O_{ij},$$
$$Q_{ij} = q_{ij}\phi_2(c_e)O_{ij},$$

for some constants $p_{ij}$ and $q_{ij}$, which determine how well each STIM binds to the Orai dimers. Each $p_{ij}$ and $q_{ij}$ is a combination of the equilibrium binding constant of the relevant Orai and STIM and the ratio of $S_1^T/S_2^T$.

The ratios of the equilibrium binding constants are estimated from the data in Fig. 3, panels M and Q. This gives $\frac{p_{11}}{p_{22}} = 1/2$, $\frac{p_{11}}{p_{33}} = 2/3$, $\frac{q_{11}}{q_{22}} = 4/5$, $\frac{q_{11}}{q_{33}} = 2$. In other words, STIM1 binds Orai2 twice as well as it binds Orai1, and binds Orai3 with an intermediate efficacy. Similarly STIM2 binds Orai1 twice as well as it binds Orai3, and binds Orai2 better still. For want of a better choice, each binding constant $p_{ij}$ with $i \neq j$ was set to $(p_{ii} + p_{jj})/2$. The relative sizes of $p_{ij}$ and $q_{ij}$ are then adjusted by choice of $S_1^T/S_2^T$. We chose $\frac{S_1^T}{S_2^T} = 3$, in order to get qualitative agreement with the data presented here.

The inward Ca current, $J_{in}$ , is given by

$$J_{\text{in}} = \alpha_0 + \sum (\alpha_{ij} P_{ij} + \beta_{ij} Q_{ij}) = \alpha_0 + \phi_1 \sum \alpha_{ij} p_{ij} O_{ij} + \phi_2 \sum \beta_{ij} q_{ij} O_{ij}.$$

for some constants $\alpha_{ij}$ and $\beta_{ij}$ which control the amount of currrent through each open STIM-Oraii,j complex. The $\alpha_{ij}$ were taken from Yoast et al (2020), and the $\beta_{ij}$ were chosen to obtain qualitative agreement with the data presented here. Reasonable agreement was obtained by setting $\beta_{ij} = \alpha_{ij}/5$, with the exceptions $\beta_{22} = 10\alpha_{22}$ and $\beta_{33} = 10\alpha_{33}$. Hence, the current through STIM2-Oraii,j is generally only 20% of the current through the analogous STIM1-Oraii,j, with the exception of Orai2,2 and Orai3,3. The parameter $\alpha_0$ is the STIM-independent leak.

We also have a conservation equation for Orai1,

$$2(O_{11} + P_{11} + Q_{11}) + O_{12} + P_{12} + Q_{12} + O_{13} + P_{13} + Q_{13} + O_1 = I_1,$$

with analogous conservation equations for Orai2 and Orai3. Given known values of $c_e$, $p_{ij}$, $q_{ij}$, $\alpha_{ij}$, $\beta_{ij}$ the conservation equations can be solved to determine $O_{ij}$, and thus determine $J_{in}$. This needs to be done for every time step of the differential equations for the calcium concentration, which are taken from [10].

To model the inhibition of the $IP_3R$ by unactivated STIM, we multiply the usual $IP_3R$ current [10] by the factor

$$F = 1 + 2\frac{K^n}{K^n + (1 - S_1)^n}\frac{K^n}{K^n + (1 - S_2)^n}$$

where K=0.005 and n=6, and where $S_i$ denotes the normalised concentration of activated STIMi. Under basal conditions, $S_1$ and $S_2$ are much less than 1, and F is thus close to 1. As $S_1$ and $S_2$ increase, F also increases, to a maximum value of 3. Thus, in the absence of any unactivated STIM, the current through the $IP_3R$ is three times greater than the current would be in the absence of feedback from STIM.

**Statistics**

All statistical tests were conducted using GraphPad Prism 8 and data presented as mean ± SEM. An unpaired, nonparametric Mann-Whitney t-test was used when comparing two sample groups. For comparisons of greater than two groups, the Kruskal-Wallis test with Dunn's multiple comparison was used. For representation of statistical significance, *, **, ***, and **** indicate p-values of $< 0.05$, $< 0.01$, $< 0.001$, and $<0.0001$ respectively.

### Legends to Supplementary Figures

**Supplementary Figure 1. Expression of all three $IP_3R$ isoforms and representative $Ca^{2+}$ oscillation profiles in WT HEK293 and STIM-KO cells.** Traces representing the three oscillatory profiles quantified throughout the manuscript. (A) Non-responding cells demonstrate no response or a single initial spike that returns to baseline following agonist stimulation. (B) Cells showing regenerative oscillations that return to baseline prior to each subsequent spike for the duration of the recording. (C) Plateau cells with sustained elevation of cytosolic $Ca^{2+}$ for the duration of the recording. The arrow in each trace represents the time of addition of 10μM carbachol (CCh). (D) Western blot analysis of $IP_3R1/2/3$ isoforms in WT HEK293 and STIM knockout cell lines. (E) Quantification of $IP_3R1$ intensity normalized to α-tubulin loading control. Each point represents an individual biological replicate (*n* = 3 each for each condition). (F) Quantification of $IP_3R2$ intensity (*n* = 4 each for each condition). (G) Quantification of $IP_3R3$ intensity (*n* = 3 each for each condition). (H) Characteristic $Ca^{2+}$ oscillation profiles in WT HEK293, (I) STIM1-KO, (J) STIM2-KO, and (K) STIM1/2-KO cell lines in response to 10μM CCh stimulation (represented by arrow in H).

**Supplementary Figure 2. Rapid oscillatory profile of HEK293 STIM1/2-KO cells is not due STIM-independent activation of Orai channels.** (A) $Ca^{2+}$ traces from 5 representative cells/condition upon stimulation with 10 μM carbachol (CCh) at 1 minute (indicated by arrow in A) in the presence of 2mM external $Ca^{2+}$. WT HEK293 cells were stimulated in the absence or (C) presence of 5μM $Gd^{3+}$. (B) Representative $Ca^{2+}$ traces of STIM1/2-KO cells in the absence or (D) presence of 5μM $Gd^{3+}$. (E) Quantification of total oscillations in 14 minutes from (A-D). From left to right *n* = 189, 90, 78, and 113 individual cells. Scatter plot in (E) is presented as mean ± SEM and analyzed with the Kruskal-Wallis one-way ANOVA with multiple comparisons to WT HEK293 cells (*p<0.05; ****p<0.0001; ns, not significant). Comparison between STIM1/2-KO cells in the absence or presence of 5μM $Gd^{3+}$ was performed with the Mann-Whitney U test (ns, not significant).

**Supplementary Figure 3. $Ca^{2+}$ oscillation data from two independent clones for each of HEK293 STIM1-KO, STIM2-KO, and STIM1/2-KO cells.** (A) WT HEK293 $Ca^{2+}$ traces from 5 representative cells/condition upon stimulation with 10 μM carbachol (CCh) at 1 minute (indicated by arrow in A) in the presence of 2mM external $Ca^{2+}$. Same conditions as in (A) but for (B) STIM1-KO clones# 1 and 4, (C) STIM2-KO clones# 6 and 14, and (D) STIM1/2-KO clones# 1 and 20. (E) Quantification of total oscillations in 14 minutes. From left to right *n* = 189, 120, 119, 150, 175, 86, and 62 individual cells. Oscillation frequency data in (Fig. 1I) is pooled from the two clones presented in (B-D) for each condition. (F) % oscillating cells, (G) % plateau cells, and (H) % non-responding cells for each cell line shown in (A-D). For (F-H) from left to right *n* = 6, 3, 3, 6, 6, 3 and 3 independent experiments. Scatter plots of oscillation frequency data (E) are presented as mean ± SEM and analyzed with the Kruskal-Wallis one-way ANOVA with multiple comparisons to WT HEK293 cells (****p<0.0001; ns, not significant). Scatter plots in (F-H) are presented as mean ± SEM and analyzed with an ordinary one-way ANOVA with multiple comparisons to WT HEK293 cells (*p<0.05; **p<0.01; ***p<0.001; ****p<0.0001; ns, not significant).

**Supplementary Figure 4. Western blot analysis of tk-STIM constructs, double Orai-STIM knockout cell lines, and CRAC currents mediated by STIM chimera constructs.** (A) *Left:* Representative western blots of WT HEK293, STIM1/2-KO cells, and STIM1/2-KO cells rescued with tk-YFP-STIM1 or tk-YFP-STIM2 and probed with primary antibodies against STIM1 and GAPDH. *Right*: Same samples shown in *Left*, but probed with a primary antibody against STIM2. (B) Representative western blots of double Orai-KO/single STIM-KO HEK293 cells probed with primary antibodies against STIM1, STIM2, and GAPDH. (C) Representative I/V curves of whole-cell patch clamp currents in STIM1/2-KO cells expressing CFP-Orai1 with either YFP-S1N-S2C or YFP-S2N-S1C chimeric STIM constructs. In these recordings, store depletion was achieved through dialysis of 20mM BAPTA through the patch pipette. (D) Quantification of peak current density at -100mV from (C). For each condition *n* = 8 (S1N-S2C + Orai1) and *n* = 7 (S2N-S1C + Orai1) individual cells. Scatter plot in (D) is presented as mean ± SEM and analyzed with the Mann-Whitney U test (***p<0.001).

**Supplementary Figure 5. Quantifications of $Ca^{2+}$ signaling events in STIM-KO cells with the lanthanide insulation protocol or rescue with the non-agonist STIM mutants.** Quantification of (A) % oscillating cells, (B) % plateau cells, and (C) % non-responding cells for each cell type shown in (Fig. 4A-D). From left to right *n* = 6, 6, 6, and 6 independent experiments. (D) Representative $Ca^{2+}$ oscillation traces of STIM1/2-KO cells and (E) STIM1/2-KO cells rescued with combined YFP-STIM2.1 + STIM1-F394H-CFP and stimulated with 10 µM carbachol (CCh) at 1 minute (indicated by arrow in D) in the presence of 1mM $Gd^{3+}$ and 2mM external $Ca^{2+}$. (F) Quantification of total oscillations in 14 minutes from (D-E). From left to right *n* = 62 (STIM1/2-KO) and *n* = 77 (STIM2.1+STIM1-F394H) individual cells. (G) % oscillating cells, (H) % plateau cells, and (I) % non-responding cells for each cell type shown in (D-E). For (G-I) from left to right *n* = 3 independent experiments for both conditions. Quantification of (J) % oscillating cells, (K) % plateau cells, and (L) % non-responding cells for each cell line shown in (Fig. 4K-O). From left to right *n* = 5, 3, 3, and 3 independent experiments. Quantification of (M) % oscillating cells, (N) % plateau cells, and (N) % non-responding cells for each cell line shown in (Fig. 4R-V). From left to right *n* = 3, 3, 3, and 4 independent experiments. Scatter plots in (A-C, J-L, M-O) are presented as mean ± SEM and analyzed with an ordinary one-way ANOVA with multiple comparisons to WT HEK293 (A-C) or STIM1/2-KO (J-L, M-O) where (*p<0.05; ***p<0.001; ns, not significant). Scatter plots in (F-I) are presented as mean ± SEM and analyzed with an unpaired Student's t-test (F) or Mann-Whitney U test (*p<0.05; ns, not significant).

**Supplementary Figure 6. HEK293 STIM-KO cells show similar rates of ER $Ca^{2+}$ depletion in response to SERCA inhibition.** (A) Measurement of ER $Ca^{2+}$ in HEK293 STIM-KO cells using R-CEPIA1*er*. Cells were stimulated with 25µM CPA in a 0mM $Ca^{2+}$ bath solution followed by re-addition of 2mM $Ca^{2+}$ at 15 minutes (without CPA). Inset shows zoomed in trace from 3-9 minutes (B) Quantification of ER $Ca^{2+}$ depletion rate in (A) at 5 minutes after CPA stimulation in 0mM external $Ca^{2+}$. (C) Quantification of ER $Ca^{2+}$ refilling rate in (A) at 20 minutes after restoration of 2mM external $Ca^{2+}$. From left to right *n* = 63, 56, 57, and 70 individual cells. Scatter plots of ER $Ca^{2+}$ depletion and refilling quantifications are presented as mean ± SEM and analyzed with the Kruskal-Wallis one-way ANOVA with multiple comparisons to WT HEK293 (****p<0.0001; ns, not significant).

**Supplementary Figure 7. STIM1/STIM2 regulate IP3R-mediated $Ca^{2+}$ puffs in response to carbachol stimulation.** $Ca^{2+}$ puffs were elicited by addition of 1 μM CCh to the bath in WT-HEK293 (n= 9 independent experiments with 63 cells) and STIM1/2-KO cells (n= 5 independent experiments with 65 cells) and images were acquired at a frame rate of 91 (fps). (A) The number of puffs were significantly higher in STIM1/2-KO (227 in 65 cells) when compared to HEK293 cells (164 in 63 cells). Similarly, (B) The number of puff sites were significantly higher in STIM1/2-KO (118 in 65 cells) when compared to HEK293 cells (92 in 63 cells). (C) Bar graph showing the number of WT-HEK293 and STIM1/2-KO cells in which the calcium signals globalize within 0-60 seconds or 61-90 seconds following CCh perfusion. Also, shown are the number of cells in which the calcium signals do not globalize. (D) Mean rise and decay times for the fluorescence to increase or decrease to 20%, 50%, 80%, 100% of $Ca^{2+}$ puffs evoked by CCh from HEK293 (n=63) and STIM1/2-KO (n=65) cells are plotted. The mean rise and decay times for the $Ca^{2+}$ puffs did not differ between HEK-293 and STIM1/2-KO. (E) Histogram representing the amplitudes distribution of the $Ca^{2+}$ puffs following stimulation with CCh of WT-HEK293 and STIM1/2-KO cells. The amplitudes distribution of the $Ca^{2+}$ puffs did not differ between HEK293 and STIM1/2-KO cells. Data presented as mean ± SEM. Statistical significance was determined by student's t-test (unpaired, two-tailed). *p<0.05, **p<0.01 with respect to HEK293. ns, not significant.

**Supplementary Figure 8. Mathematical model simulations of $Ca^{2+}$ responses to increasing agonist concentrations under the $Gd^{3+}$ insulation protocol in WT and STIM-KO cells.** Responses to four different agonist concentrations ($p$ = 0.015, $p$ = 0.03, $p$ = 0.06, $p$ = 0.25) applied sequentially in that order, each for 5 min. In each panel the solid line is the $Ca^{2+}$ concentration, and is plotted against the left axis, while the dotted and dashed lines are the STIM-mediated currents, and are plotted against the right axis. Panels A-D; $Ca^{2+}$ responses in a closed cell system, mimicking the 1mM $Gd^{3+}$ insulation protocol (i.e., with no membrane influx of extrusion of $Ca^{2+}$) with all three Orai isoforms present. The STIM1/2-KO cell is the most effective oscillator, which results from the model assumption that the IP3R is inhibited by unactivated STIM. Under this $Gd^{3+}$ insulation protocol (closed cell), the absence of STIM1 (B and D) does not hinder the cell from responding with a raised plateau of $Ca^{2+}$ at high agonist concentrations.

**Supplementary Figure 9. Mathematical model simulations of $Ca^{2+}$ responses to thapsigargin and increasing agonist concentrations in double Orai-KO/single STIM-KO cells.** Responses to thapsigargin (panels A1, B1 and C1) and responses to increasing steps of agonist concentration ($p$ = 0.015, $p$ = 0.03, $p$ = 0.06, $p$ = 0.25) applied sequentially in that order, each for 5 min (other panels). The left panels (A1, A2, A3, A4) are for an Orai2,3-DKO cell, the middle panels (B1, B2, B3, B4) are for an Orai2,3-DKO cell, and the right panels (C1, C2, C3,

C4) are for an Orai1,3-DKO cell. For each column of panels, the color coding described in the top panel refers also to the lower panels. These responses reproduce the qualitative features seen in the experimental data (**Fig. 3**). As seen also in **Fig. 5** and **S7**, cells are unable to maintain a raised $Ca^{2+}$ plateaus in the absence of STIM1.

**Supplementary Figure 10. Mathematical model simulations of $Ca^{2+}$ influx (net current) mediated by STIM1 or STIM2 interactions with various Orai homomers and heteromers.** Simulations of STIM1-mediated (panel A) and STIM2-mediated (panel B) $Ca^{2+}$ currents in response to increasing concentrations of agonist (denoted by p) in various Oraij,k conditions.

**Supplementary video 1 and 2.**

Images in Cal520 loaded WT-HEK293 (video 1) or STIM1/2-KO cells (video 2) were captured at 166 fps in TIRF mode as described in Methods. Movies shows 3 seconds prior to photolysis of ci$IP_3$ (caged $IP_3$) and the following 60 seconds of activity. Photolysis was accomplished by exposure from a 405 nm laser for 1 second (indicated by the yellow circle). ci$IP_3$; 6-*O*-[(4,5-Dimethoxy-2-nitrophenyl)methyl]-2,3-*O*-(1-methylethylidene)-D-*myo*-Inositol 1,4,5-tris[bis[(1-oxopropoxy)methyl]phosphate]

# S1

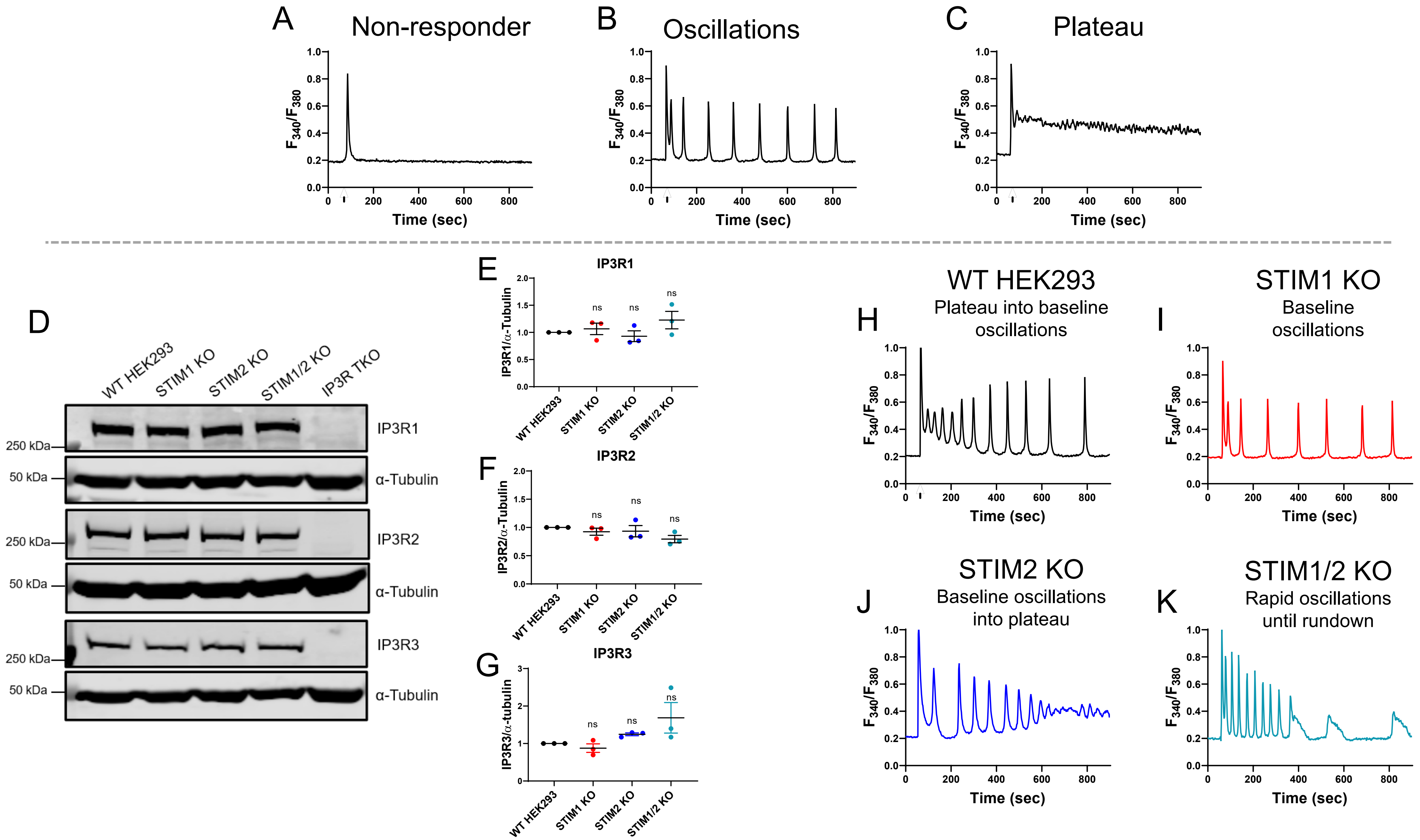

A
Non-responder
B
Oscillations
C
Plateau
F340/F380
Time (sec)
D
WT HEK293
STIM1 KO
STIM2 KO
STIM1/2 KO
IP3R TKO
IP3R1
α-Tubulin
IP3R2
IP3R3
250 kDa
50 kDa
E
IP3R1
IP3R1/α-Tubulin
F
IP3R2
IP3R2/α-Tubulin
G
IP3R3
IP3R3/α-tubulin
ns
H
WT HEK293
Plateau into baseline oscillations
I
STIM1 KO
Baseline oscillations
J
STIM2 KO
Baseline oscillations into plateau
K
STIM1/2 KO
Rapid oscillations until rundown

S2

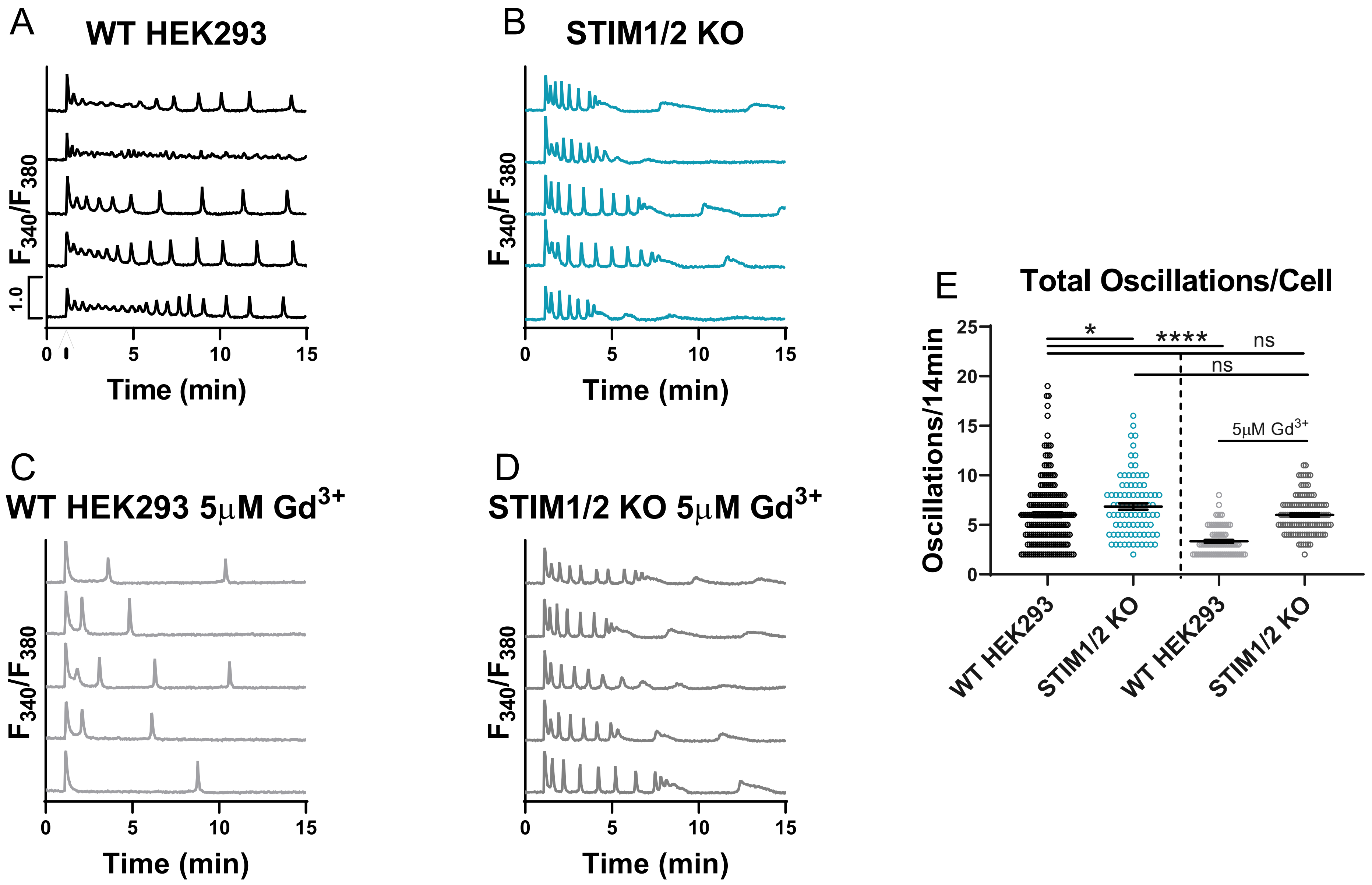
A
WT HEK293
$F_{340}/F_{380}$
1.0
Time (min)
B
STIM1/2 KO
$F_{340}/F_{380}$
Time (min)
C
WT HEK293 5μM $Gd^{3+}$
$F_{340}/F_{380}$
Time (min)
D
STIM1/2 KO 5μM $Gd^{3+}$
$F_{340}/F_{380}$
Time (min)
E
Total Oscillations/Cell
Oscillations/14min
*
****
ns
ns
5μM $Gd^{3+}$
WT HEK293
STIM1/2 KO
WT HEK293
STIM1/2 KO

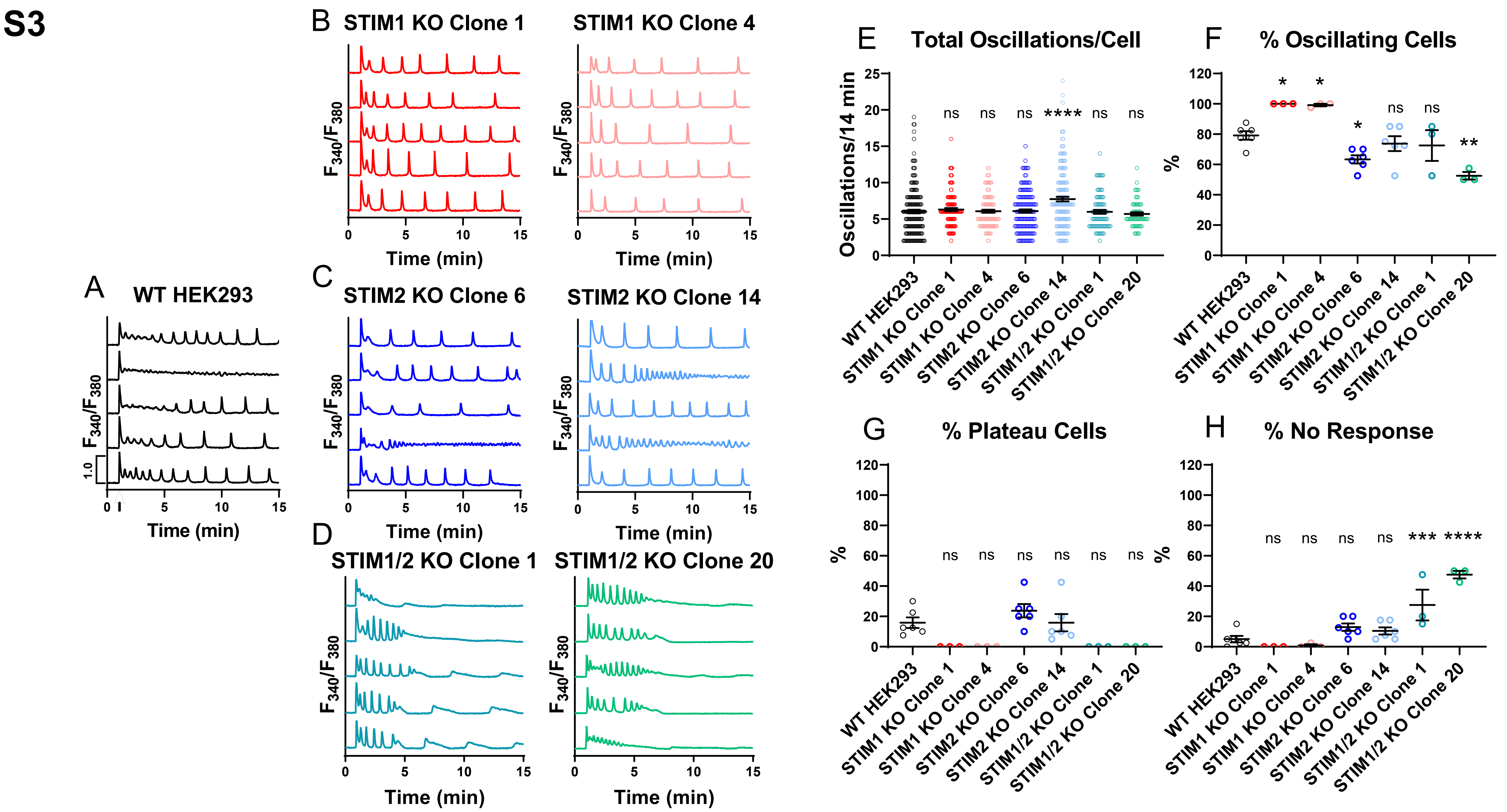
S3
A WT HEK293
B STIM1 KO Clone 1
STIM1 KO Clone 4
C STIM2 KO Clone 6
STIM2 KO Clone 14
D STIM1/2 KO Clone 1
STIM1/2 KO Clone 20
F340/F380
Time (min)
E Total Oscillations/Cell
Oscillations/14 min
F % Oscillating Cells
G % Plateau Cells
H % No Response
%
WT HEK293
STIM1 KO Clone 1
STIM1 KO Clone 4
STIM2 KO Clone 6
STIM2 KO Clone 14
STIM1/2 KO Clone 1
STIM1/2 KO Clone 20
ns
*
**
***
****

**S4**

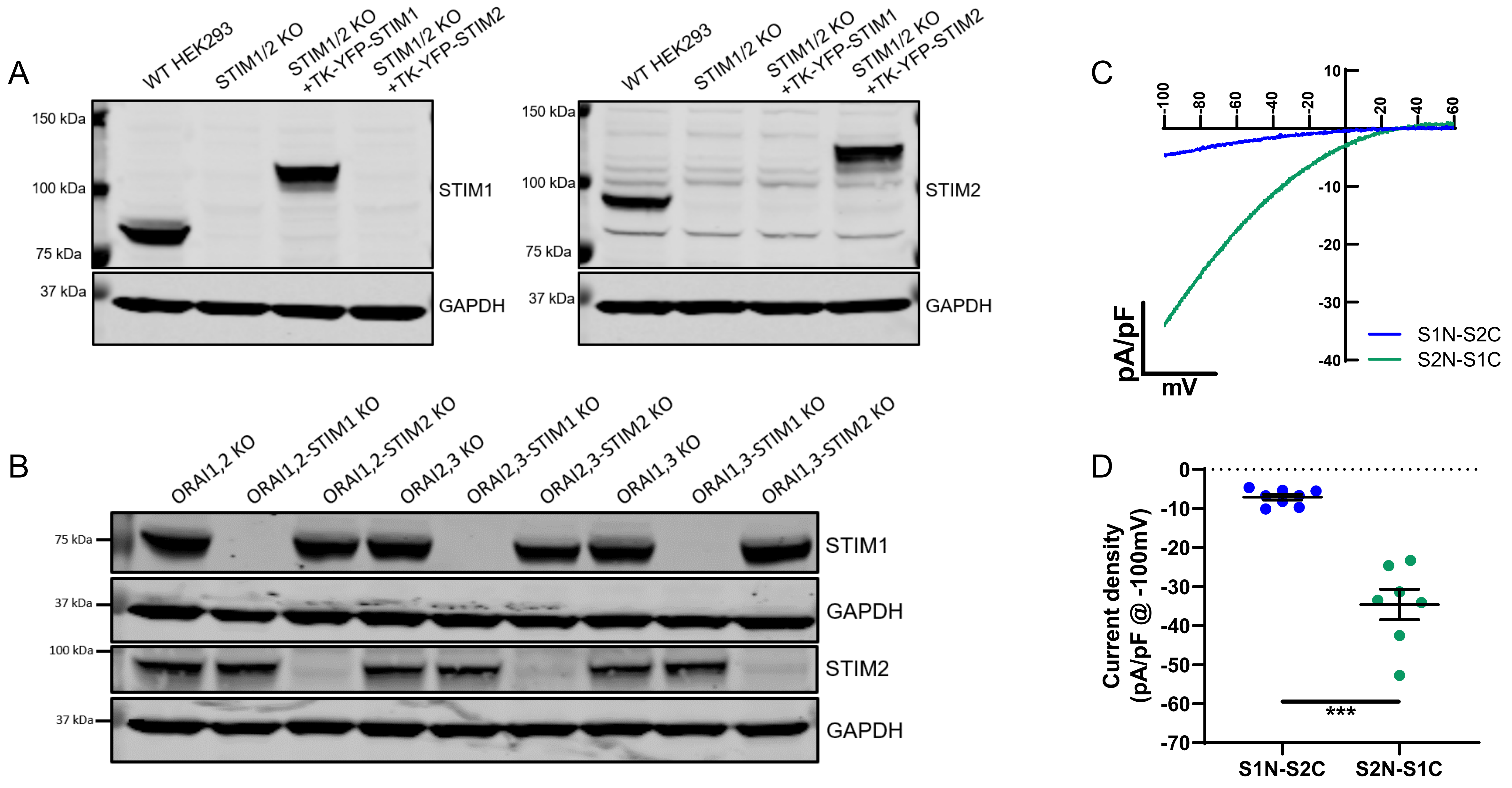

A
WT HEK293
STIM1/2 KO
STIM1/2 KO +TK-YFP-STIM1
STIM1/2 KO +TK-YFP-STIM2
150 kDa
100 kDa
75 kDa
37 kDa
STIM1
GAPDH
STIM2
B
ORAI1,2 KO
ORAI1,2-STIM1 KO
ORAI1,2-STIM2 KO
ORAI2,3 KO
ORAI2,3-STIM1 KO
ORAI2,3-STIM2 KO
ORAI1,3 KO
ORAI1,3-STIM1 KO
ORAI1,3-STIM2 KO
C
pA/pF
mV
S1N-S2C
S2N-S1C
D
Current density (pA/pF @ -100mV)
***

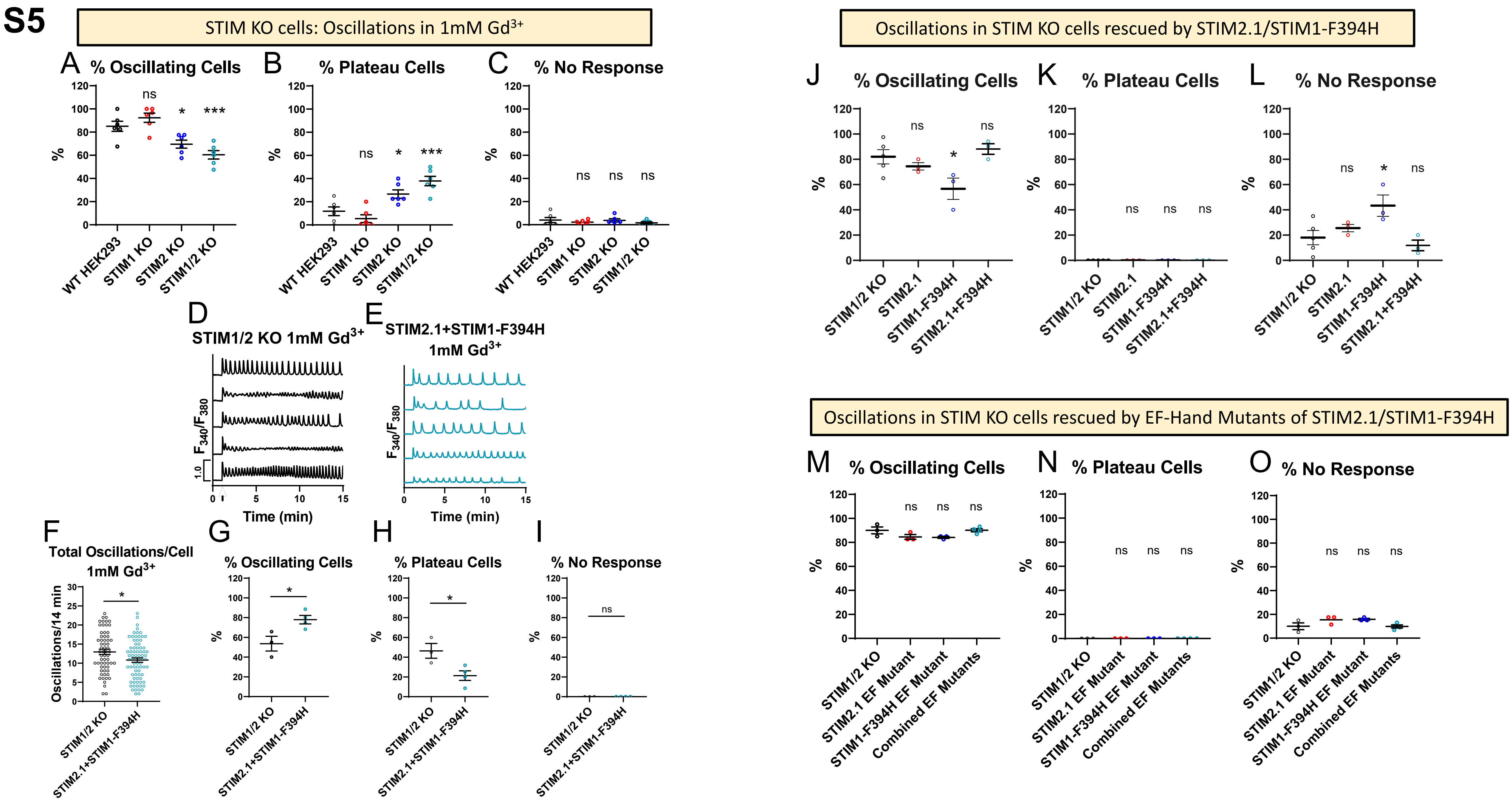

S5
STIM KO cells: Oscillations in 1mM Gd3+
A % Oscillating Cells
B % Plateau Cells
C % No Response
WT HEK293
STIM1 KO
STIM2 KO
STIM1/2 KO
D STIM1/2 KO 1mM Gd3+
E STIM2.1+STIM1-F394H 1mM Gd3+
F340/F380
Time (min)
F Total Oscillations/Cell 1mM Gd3+
Oscillations/14 min
G % Oscillating Cells
H % Plateau Cells
I % No Response
STIM2.1+STIM1-F394H
Oscillations in STIM KO cells rescued by STIM2.1/STIM1-F394H
J % Oscillating Cells
K % Plateau Cells
L % No Response
STIM2.1
STIM1-F394H
STIM2.1+F394H
Oscillations in STIM KO cells rescued by EF-Hand Mutants of STIM2.1/STIM1-F394H
M % Oscillating Cells
N % Plateau Cells
O % No Response
STIM2.1 EF Mutant
STIM1-F394H EF Mutant
Combined EF Mutants

**S6**

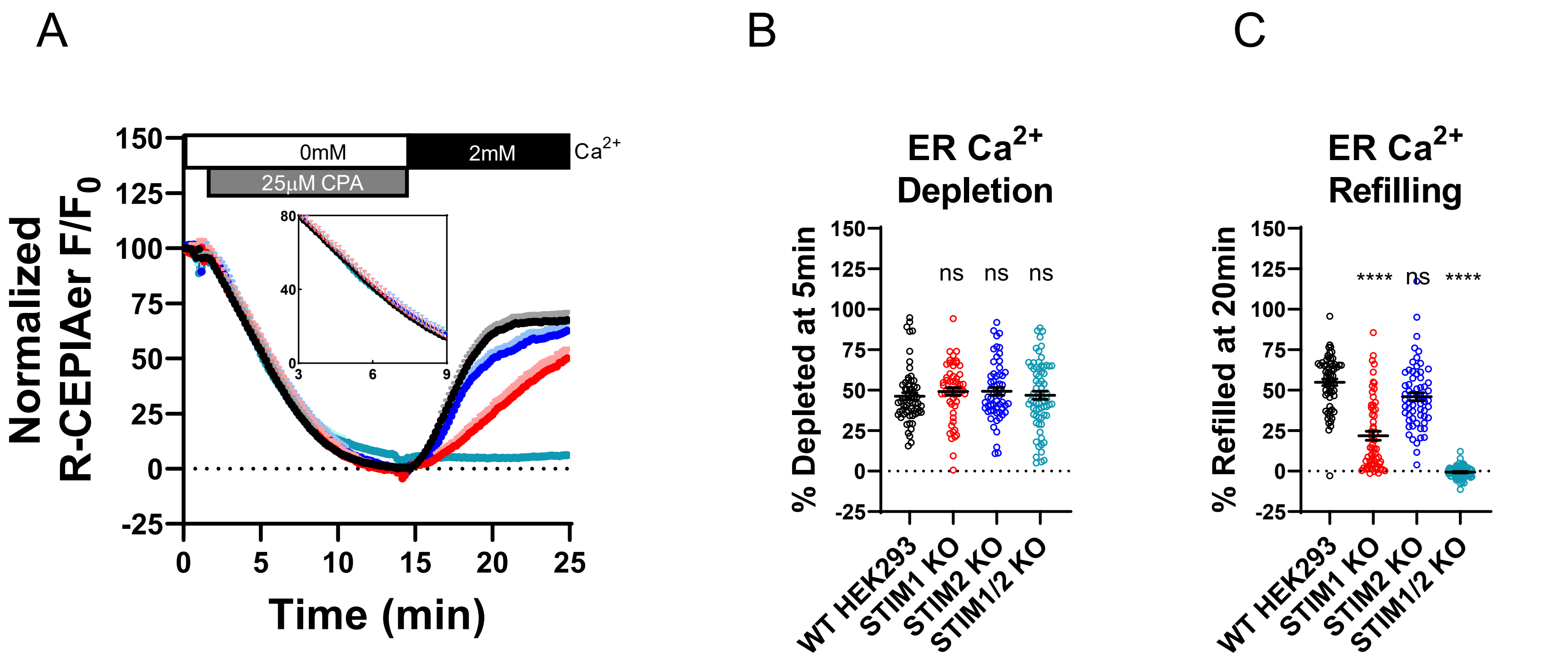

A
B
C
150
125
100
75
50
25
0
-25
0mM
2mM
Ca2+
25µM CPA
80
40
0
3
6
9
Normalized R-CEPIAer F/F0
0
5
10
15
20
25
Time (min)
ER Ca2+ Depletion
% Depleted at 5min
ns
ns
ns
WT HEK293
STIM1 KO
STIM2 KO
STIM1/2 KO
ER Ca2+ Refilling
% Refilled at 20min
****
ns
****
WT HEK293
STIM1 KO
STIM2 KO
STIM1/2 KO

S7

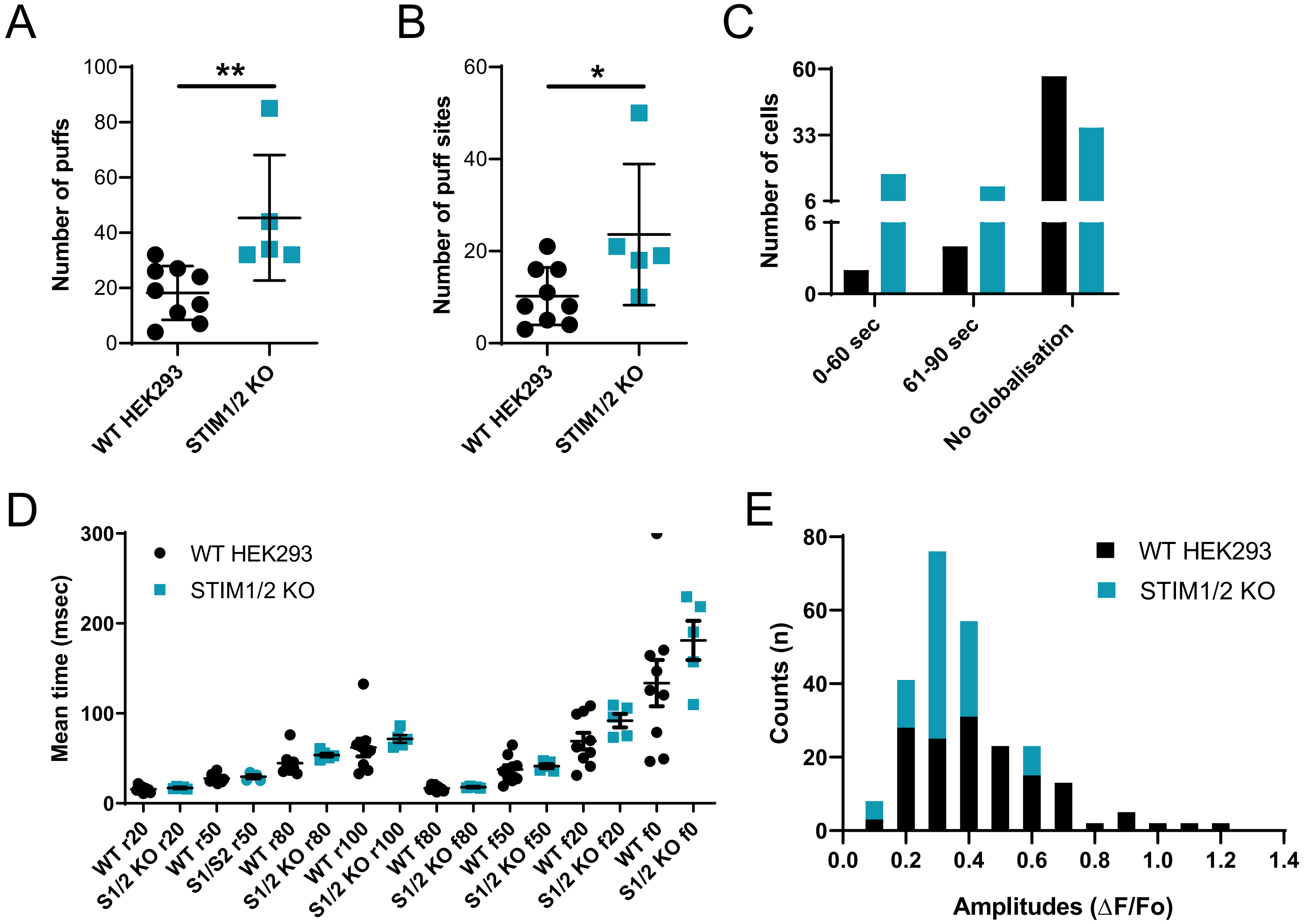
A
Number of puffs
**
WT HEK293
STIM1/2 KO
B
Number of puff sites
*
WT HEK293
STIM1/2 KO
C
Number of cells
0-60 sec
61-90 sec
No Globalisation
D
Mean time (msec)
WT HEK293
STIM1/2 KO
WT r20
S1/2 KO r20
WT r50
S1/S2 r50
WT r80
S1/2 KO r80
WT r100
S1/2 KO r100
WT f80
S1/2 KO f80
WT f50
S1/2 KO f50
WT f20
S1/2 KO f20
WT f0
S1/2 KO f0
E
Counts (n)
WT HEK293
STIM1/2 KO
Amplitudes (ΔF/Fo)

# S8

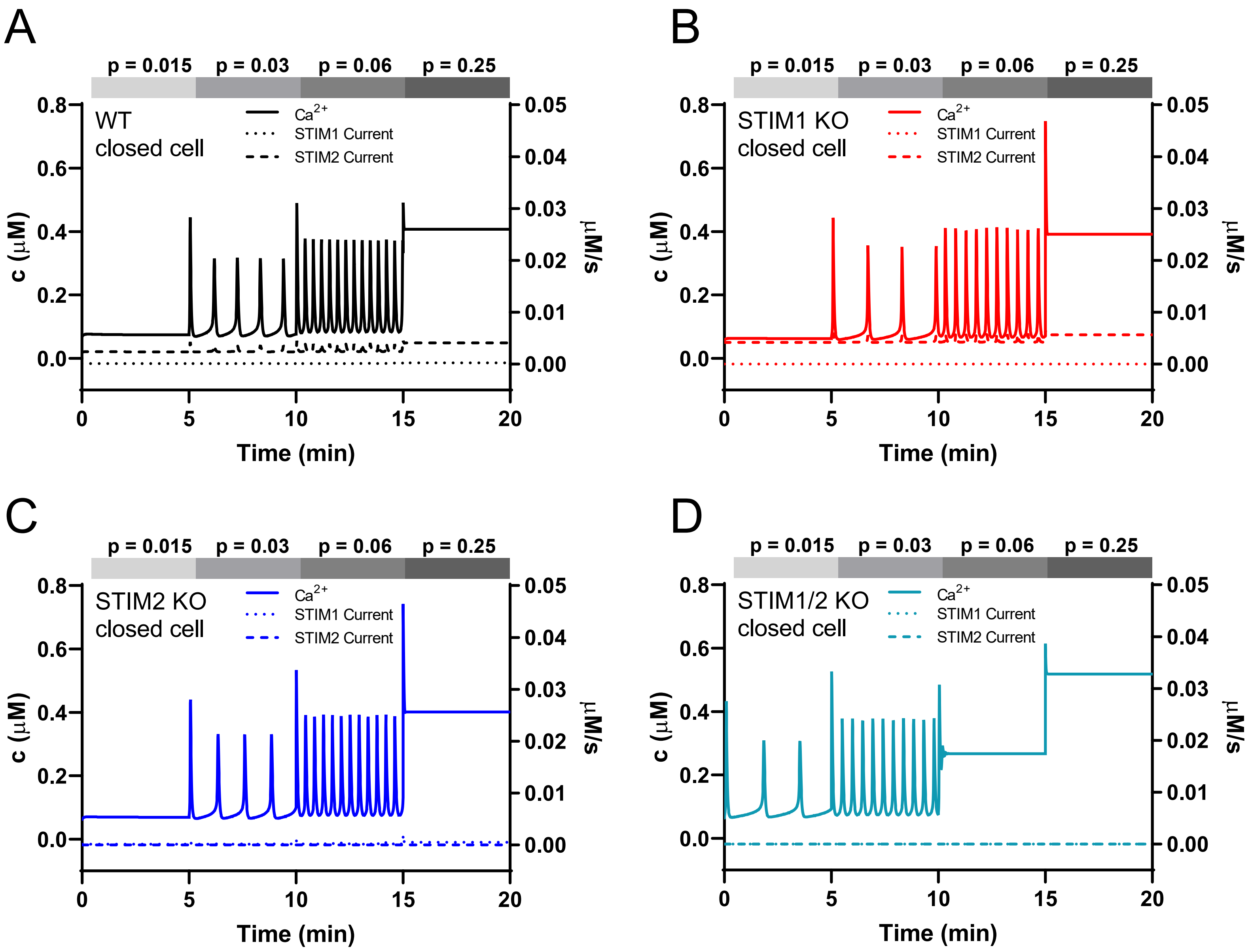

A
p = 0.015
p = 0.03
p = 0.06
p = 0.25
WT
closed cell
Ca2+
STIM1 Current
STIM2 Current
c (μM)
μM/s
Time (min)
B
STIM1 KO
closed cell
C
STIM2 KO
closed cell
D
STIM1/2 KO
closed cell

**S9**

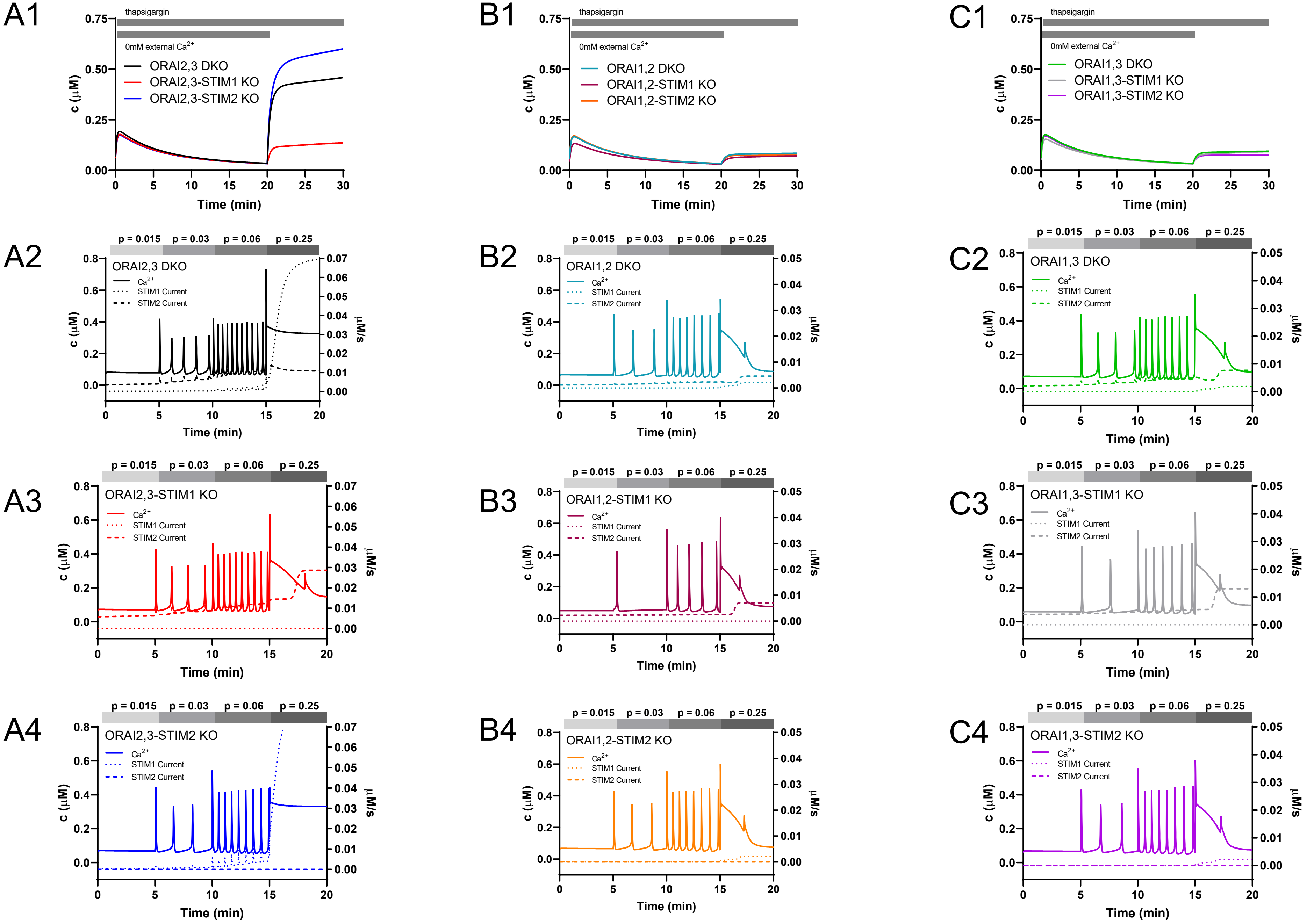
A1
thapsigargin
0mM external $Ca^{2+}$
ORAI2,3 DKO
ORAI2,3-STIM1 KO
ORAI2,3-STIM2 KO
c (μM)
Time (min)
B1
thapsigargin
0mM external $Ca^{2+}$
ORAI1,2 DKO
ORAI1,2-STIM1 KO
ORAI1,2-STIM2 KO
c (μM)
Time (min)
C1
thapsigargin
0mM external $Ca^{2+}$
ORAI1,3 DKO
ORAI1,3-STIM1 KO
ORAI1,3-STIM2 KO
c (μM)
Time (min)
A2
p = 0.015 p = 0.03 p = 0.06 p = 0.25
ORAI2,3 DKO
$Ca^{2+}$
STIM1 Current
STIM2 Current
c (μM)
μM/s
Time (min)
B2
p = 0.015 p = 0.03 p = 0.06 p = 0.25
ORAI1,2 DKO
$Ca^{2+}$
STIM1 Current
STIM2 Current
c (μM)
μM/s
Time (min)
C2
p = 0.015 p = 0.03 p = 0.06 p = 0.25
ORAI1,3 DKO
$Ca^{2+}$
STIM1 Current
STIM2 Current
c (μM)
μM/s
Time (min)
A3
p = 0.015 p = 0.03 p = 0.06 p = 0.25
ORAI2,3-STIM1 KO
$Ca^{2+}$
STIM1 Current
STIM2 Current
c (μM)
μM/s
Time (min)
B3
p = 0.015 p = 0.03 p = 0.06 p = 0.25
ORAI1,2-STIM1 KO
$Ca^{2+}$
STIM1 Current
STIM2 Current
c (μM)
μM/s
Time (min)
C3
p = 0.015 p = 0.03 p = 0.06 p = 0.25
ORAI1,3-STIM1 KO
$Ca^{2+}$
STIM1 Current
STIM2 Current
c (μM)
μM/s
Time (min)
A4
p = 0.015 p = 0.03 p = 0.06 p = 0.25
ORAI2,3-STIM2 KO
$Ca^{2+}$
STIM1 Current
STIM2 Current
c (μM)
μM/s
Time (min)
B4
p = 0.015 p = 0.03 p = 0.06 p = 0.25
ORAI1,2-STIM2 KO
$Ca^{2+}$
STIM1 Current
STIM2 Current
c (μM)
μM/s
Time (min)
C4
p = 0.015 p = 0.03 p = 0.06 p = 0.25
ORAI1,3-STIM2 KO
$Ca^{2+}$
STIM1 Current
STIM2 Current
c (μM)
μM/s
Time (min)

S10

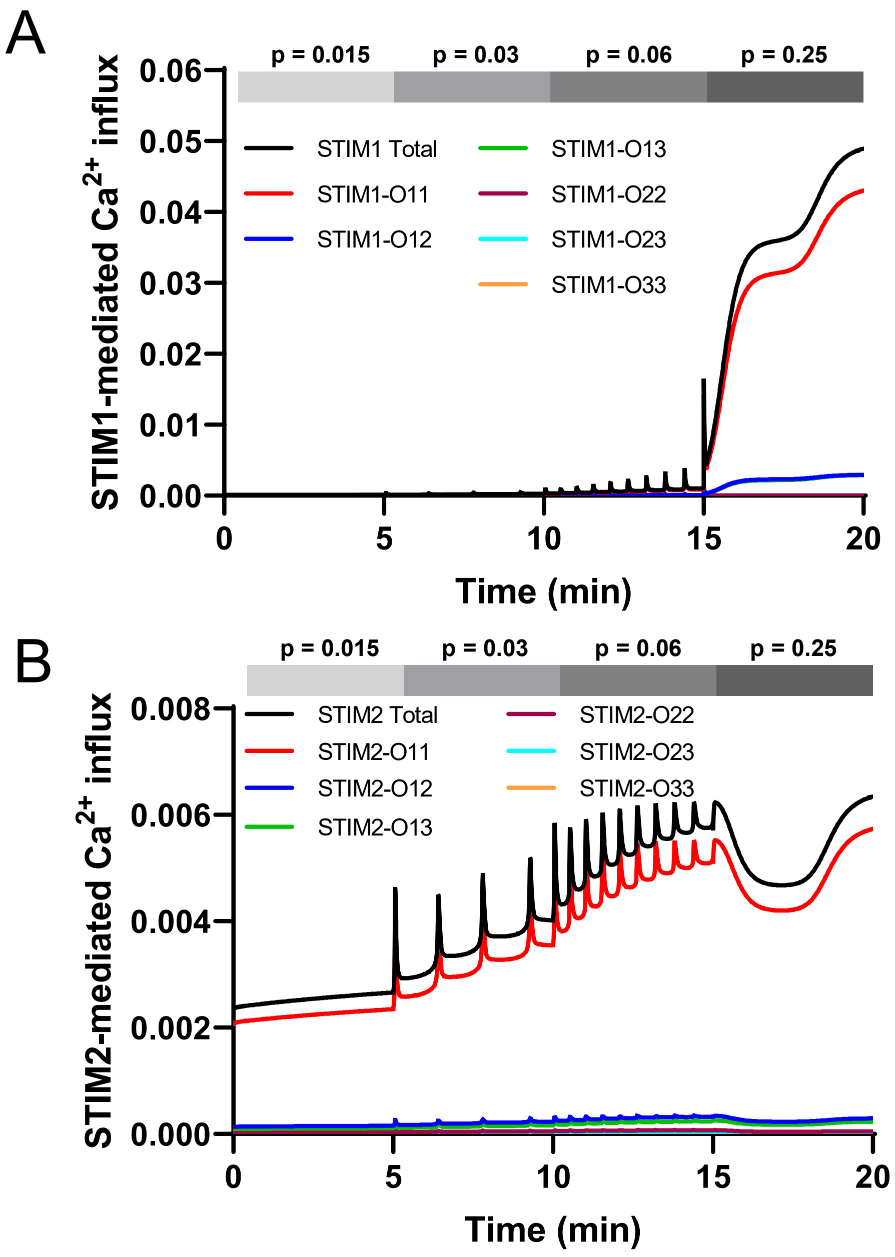

A
p = 0.015
p = 0.03
p = 0.06
p = 0.25
STIM1-mediated Ca2+ influx
0.06
0.05
0.04
0.03
0.02
0.01
0.00
STIM1 Total
STIM1-O11
STIM1-O12
STIM1-O13
STIM1-O22
STIM1-O23
STIM1-O33
0
5
10
15
20
Time (min)
B
p = 0.015
p = 0.03
p = 0.06
p = 0.25
STIM2-mediated Ca2+ influx
0.008
0.006
0.004
0.002
0.000
STIM2 Total
STIM2-O11
STIM2-O12
STIM2-O13
STIM2-O22
STIM2-O23
STIM2-O33
0
5
10
15
20
Time (min)
